# How differing modes of transgenerational inheritance affect population viability in fluctuating environments

**DOI:** 10.1101/506865

**Authors:** Stephen R. Proulx, Snigdhadip Dey, Thiago Guzella, Henrique Teotónio

## Abstract

Different modes of transgenerational inheritance are expected to affect population persistence in fluctuating environments. We here analyze *Caenorhabiditis elegans* density-independent per capita growth rate time series on 36 populations experiencing 6 controlled sequences of challenging oxygen level fluctuations across 60 generations, and parameterize competing models of transgenerational inheritance that do not involve genetic changes in order to explain observed dynamics. Our analysis shows that phenotypic plasticity and anticipatory maternal effects are sufficient to explain growth rate dynamics, but that a carryover model where epigenetic memory is imperfectly transmitted and might be reset at each generation is a better fit to the data. After validating the models, we use the fitted transgenerational inheritance parameters to forecast population growth rates and find that it is negatively related to the degree of environmental autocorrelation.

## 2 Introduction

In addition to change in average environmental directions, there has also been human-induced change in environmental variability, particularly a change in the temporal autocorrelation of fluctuating environments (Stocker *et al*., 2013; Huntingford *et al*., 2013), but a detailed understanding of its effects on population and species persistence is still missing (Lawson *et al*., 2015; Lande *et al*., 2003).

Evolutionary ecology theory focuses on the geometric mean of the population growth factor across the environmental conditions a population may face (Cohen, 1966), or stage-structured extensions (Tuljapurkar, 1982), as a measure of its long term viability in fluctuating environments. The geometric mean of population growth is sufficient to predict long term viability, and since it is independent of the sequence of environmental events (Proulx and Adler, 2010), usually only the mean and the variance of environmental conditions needs to be considered (Saether and Engen, 2015). However, several scenarios that are plausibly common can cause the temporal autocorrelation of fluctuating environments to affect population viability, including the presence of transgenerational inheritance of phenotypes that is not mediated by genetic changes across generations. While within-generation phenotypic adjustments rely on the perception of environmental cues available to the focal individuals (Chevin and Lande, 2015; Chevin and Haller, 2014), transgenerational inheritance is elicited by the perception of environmental information by the parents (Uller, 2008; Day and Bonduriansky, 2011; Leimar and McNamara, 2015). Differences in the temporal autocorrelation of environmental conditions are thus expected to affect population viability if transgenerational inheritance acts directly on individual phenotypes or creates non-Markovian density dependence across generations (Lande *et al*., 2003, 2006; Inchausti and Ginzburg, 2009).

The mode of transgenerational inheritance can be classified by whether changes in the distribution of phenotypes are based on the detection of the environment by individuals, the time point during the life-cycle when selection is acting, or on whether or not quantitative phenotypes are passed down over generations in an open-ended “cascading” way (Shea *et al*., 2011; McGlothlin and Galloway, 2014). Many empirical studies have shown that the transgenerational inheritance of phenotypes does not need to be DNA sequence based and thus that “epigenetic” and germline “reprogramming” developmental and/or physiological mechanisms are involved (Heard and Martienssen, 2014). The long-term mechanisms by which phenotypes are transmitted to future generations, and in particular how they are deterministically regulated, are however far less clear (Minkina and Hunter, 2018). In many studies, the effects of particular environmental events or genetic insults on later generations is only observed in small sub-populations of individuals artificially selected for the relevant phenotype, e.g. (Vastenhouw *et al*., 2006). In the best studied organisms, such as in the experimental models *Caenorhabditis elegans* or *Arabidopsis thaliana*, the effects of environmental history on gene expression may occur for up to 20 generations in a significant fraction of the progeny (Teixeira and Colot, 2010; Houri-Zeevi and Rechavi, 2017; Minkina and Hunter, 2018), but the consequences of specific sequences of environmental conditions have not been explored and thus how differing modes of transgenerational inheritance can influence population persistence remains uncertain (Charlesworth *et al*., 2017; Bonduriansky and Day, 2009).

Investigation of transgenerational inheritance usually uses combinatoric manipulations and these rapidly become unmanageable as the number of environmental transitions and generations increase. A complementary approach is to analyze growth rate time series data of (natural or experimental) populations experiencing a known sequence of environmental conditions and explicitly parameterize the possible modes of trans-generational inheritance. In this case, it is possible to focus directly on growth rates because this is a trait closely related to fitness and integrates all possible developmental and physiological mechanisms of transgenerational inheritance. With this goal in mind, we use here a replicated set of experimental populations of *C. elegans* where the key environmental condition, oxygen levels during embryogenesis and larval development (normoxia versus anoxia) were controlled to provide alternative patterns of environmental correlations across 60 generations (Figure S1). Our design minimized possible effects of within-generation density dependence by controlling the population size to a fixed number during larva-to-adult rearing.

In *C. elegans*, hyperosmotically stressed hermaphrodites accumulate glycerol and produce selfed embryos with reduced glycogen levels and survivorship when faced with anoxia during development until the first instar larval stage (Frazier and Roth, 2009). We previously found that the evolution of anticipatory maternal effects, in the form of glycogen embryo provisioning, underlies population viability in regularly alternating normoxia-anoxia conditions (Dey *et al*., 2016). In this prior work, we hypothesized that anticipatory high lag order effects, such as grand maternal effects, could sustain population viability under irregular fluctuating conditions. We measured the per capita density-independent population growth at each of the 60 generations in the experiments imposing irregular fluctuating anoxia conditions, and here use these observations to infer the presence and the mode of transgenerational inheritance without performing combinatoric fitness assays across environmental transitions. We develop competing statistical models to detect transgenerational inheritance due to the populations’ environmental history as phenotypic plasticity and anticipatory maternal and grand maternal effects, or to detect the degree to which a epigenetic state carries over to the next generation while implicitly considering germline reprogramming. After model validation, we use the inferred transgenerational inheritance parameters to forecast how any variable pattern of environmental fluctuation in oxygen levels can affect population viability in the immediate future.

## 3 Materials and Methods

### 3.1 Population maintenance

The GA250 population, which was exposed to a gradually increasing NaCl regime for 50 generations (Theologidis *et al*., 2014), and ultimately derived from hybridization of several wild inbred isolates and lab adaptation (Teotónio *et al*., 2012; Noble *et al*., 2017), served as the base population from which all the populations reported here were derived. GA250 is genetically diverse, with about one single nucleotide polymorphism every 250-500 bp across its 100 Mb genome (Noble *et al*., 2017), and where individuals reproduce exclusively by selfing because males are selected against in high salt (Theologidis *et al*., 2014). During the experiment we expect great loss of diversity simply by genetic drift and that certain genotypes might be selectively favored over others (Dey *et al*., 2016), but here we will assume that the different (possible) modes of transgenerational inheritance are similarly expressed by all genotypes.

Frozen samples of GA250 with > 10^3^ individuals were revived from cryogenic preserved −80°C stocks and reared for one generation in normoxia to allow expansion in population size needed for the derivation of new populations. Following our standard maintenance protocol (Teotónio *et al*., 2012, 2017), each population was reared in 10 9-cm Petri plates, each containing 28 mL of NGM-lite agar media overlaid with a lawn of HT115 *Escherichia coli*, the source of *ad libitum* food. The NGM-lite media was supplemented with NaCl to attain a salt concentration of 305 mM (1.78% w/v) (Theologidis *et al*., 2014). At the start of each generation, each Petri plate was seeded with 10^3^ starvation-arrested first larval staged individuals (L1), for physiological age synchronization. At 66 h ± 2 h after L1 seeding, adult hermaphrodites from all the 10 plates belonging to each population were mixed and harvested to a 15 mL Falcon tube with 5 mL isotonic M9 solution, followed by exposure to 1M KOH: 5% NaOCl bleach solution for 5 min ± 15 s, which killed individuals of all life stages but embryos (Teotónio *et al*., 2012). This was followed by three rounds of washing-off the bleach solution with M9 and transfer of 200 μL concentrate containing embryos plus the larval and adult debris to a 25 mM NaCl NGM-lite plate, devoid of *E. coli*. The Petri plates, one for each population, were then put inside 7L polycarbonate boxes with rubber-clamp-sealed lids (Anaero-Pack, Mitsubishi Inc.)(Figure S2).

Within one subset of the boxes, an anoxic hatching condition was achieved by placing two GasPak EZ sachets inside them (Becton, Dickinson and Company). Occurrence of anoxia condition was confirmed with two BBL Dry Anaerobic Indicator strips (Becton, Dickinson and Company). The second subset of boxes, which had normoxia hatching condition, lacked the sachets. Both types of boxes received paper towels soaked with 20 mL ddH2O to prevent the plates from desiccating. After 16 h ± 1 h, plates were taken out from the boxes and starvation-arrested synchronized live L1s were harvested with 3–5 mL M9 to a 15 mL Falcon tube. The pellet containing adult debris was removed after centrifugation at 200 rpm, and L1 density was estimated by counting the number of moving individuals in 5-10 5 uL M9 drops at 40x magnification of a Nikon SMZ1500 dissection scope. 6 h ± 1 h after opening the hatching boxes, M9 containing 10^3^ live L1s was pipetted to each of the 10 fresh NGM-lite plates to complete one generation.

Besides oxygen level conditions, we also manipulated whether maternal generations received a reliable or an unreliable light cue whenever offspring generations were exposed to anoxia. Populations from both treatments were paired to face the same sequence of oxygen level conditions over the 60 generations (see next section). Here we only use the light cue a nuisance parameter. *C. elegans* are sensitive to blue light (Ward *et al*., 2008). Petri plate rack holders (Starsted) were fitted with several strips of blue-light LED (Nichia NS6B083T and NSPB300B) in order for individuals inside the plates to have uniform light exposure (Figure S2). 48h after L1 seed blue light was 0.5 second flashed every 2 seconds during 12h, a period when hermaphrodites were expected to be undergoing oogenesis. A computer with dedicated software (LUCS) controlled light exposure for all plates at the same time, with plates that were not exposed to blue light being kept in a separate incubator.

### 3.2 Environmental sequences and replicate populations

Populations were exposed to 6 distinct sequences of normoxia-anoxia hatching environments for 60 generations (Figures 1 and S1). We generated sequences that followed the following criteria: (i) 30 generations in normoxia and 30 generations in anoxia [in order to match the same selective gradients of regularly fluctuating environments reported in Dey *et al*. (2016)]; (ii) probability of repeating the same oxygen level condition across maternal-offspring generations *ρ*_1_ of 0.46 (negligible autocorrelation at lag 1), in order for mothers to have little information about their offspring environment; (iii) difference between the number of consecutive generations in normoxia or anoxia not larger than 3, to even the possibility of long term effects between normoxia and anoxia; and (iv) probability of repeating the same oxygen level condition at grand maternal and grand offspring generations *ρ*_2_ of 0.52 (negligible autocorrelation at lag 2), of 0.35 (negative autocorrelation at lag 2) or of 0.62 (positive autocorrelation at lag2).

**Figure 1:**
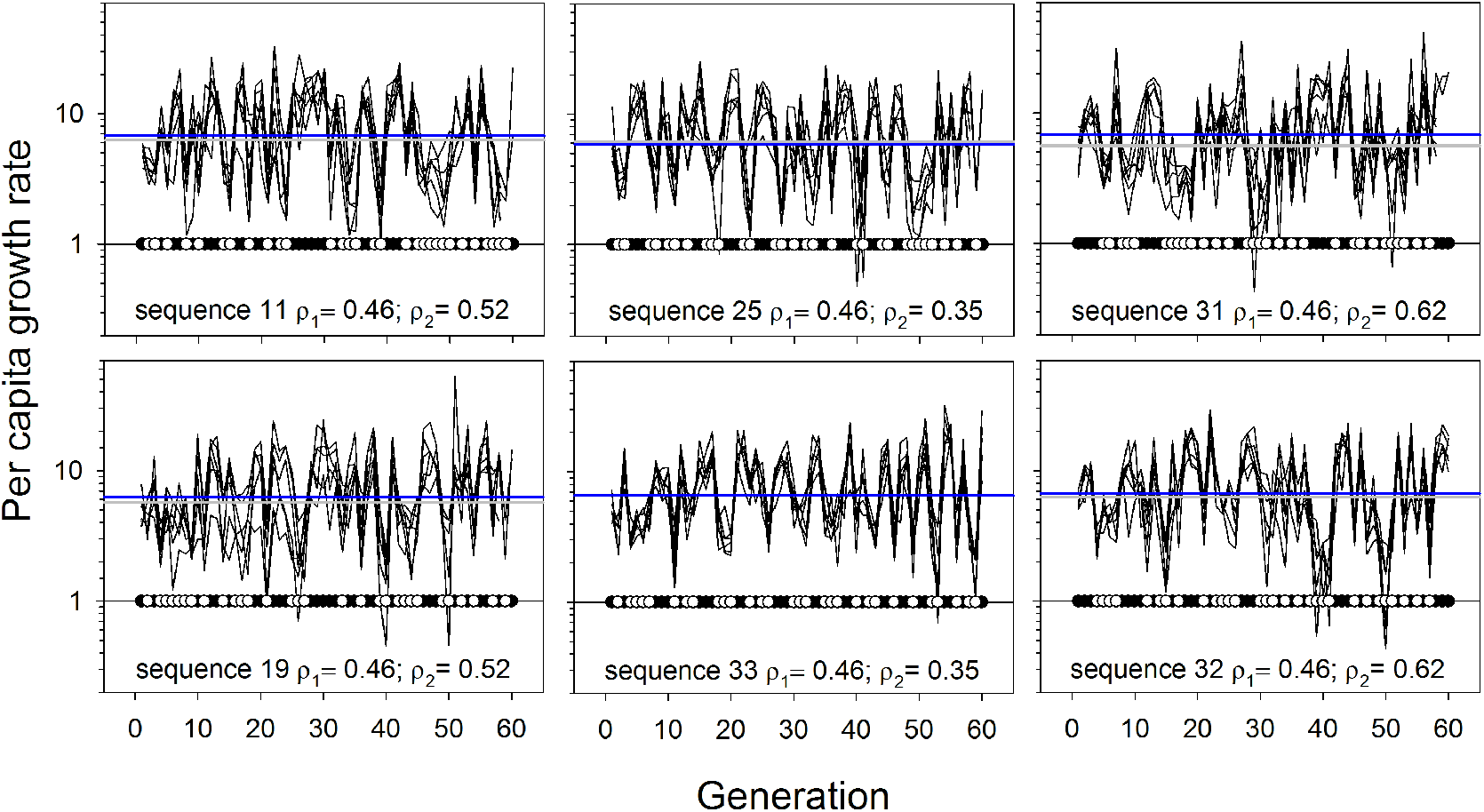
Time series of the measured per capita growth rate for 6 populations per environmental sequence (logarithmic scale). Insets show the probability of repeating the same oxygen level condition (filled dots normoxia, white dots anoxia) at lag 1 (*ρ*_1_) or lag 2 (*ρ*_2_), over 59 or 58 environmental transitions, respectively. Horizontal lines indicate the average geometric mean growth rates across 60 generations: blue for the 3 reliable populations, gray for the 3 unreliable populations.

Of the sequences retrieved, two from each of the criteria in (iv) were employed: sequence 11 (replicate populations U1, U2, and U9 for the “<u>U</u>nreliable” light cue treatment, and replicate populations R1-R3 for the “<u>R</u>eliable” light cue treatment) and sequence 19 (U3, U4, U10; R4-R6) for lag 1 and lag 2 autocorrelations of close to zero; sequence 25 (U5, U6, U11; R7-R9) and sequence 33 (U16-U18; R16-R18) for negative lag 2 autocorrelations; and sequence 31 (U7, U8, U12; R10-R12) and sequence 32 (U13-U15; R13-R15) for positive lag 2 autocorrelations (Figure S1). A total of 36 populations were thus followed (Figure 1).

### 3.3 Density-independent population growth measurements

The per-capita growth factor was calculated in each generation by dividing the live L1 density after varying oxygen level exposure by 10^4^, the number of L1s seeded in the prior generation (Figure 1). This measure is a lower bound of the per-capita density-independent population growth factor, since many live L1s are lost when removing larval and adult debris by centrifugation, before live L1 counting. For this reason, some of the observed values even if below 1 do not mean negative population growth (and we were able to seed 10^4^ L1s in each generation). The population growth factor is often used in population biology to measure the discrete time growth of a population, and the natural log of the growth factor is roughly equivalent to the instantaneous population growth rate. In our statistical treatment we use the measured growth factor to infer the parameters that determine population viability.

The experiments were run in two different “blocks”, each defined by the GA250 thawing date (2012 and 2015) and by the location where the experiments were done (Lisbon and Paris). A random id was assigned to each population to avoid potential bias during handling. The growth factor was measured by the same experimenter (SD). Data from populations U1-U8 was previously presented in Figure 3B of Dey *et al*. (2016). A total of n=2128 observations were available for analysis (Figure 1).

### 3.4 Inference of transgenerational inheritance

We used a Bayesian approach to jointly infer the direct environmental effects and the type of transgenerational inheritance that fit the observed population growth dynamics in the 60 generation experiments. For inference we used only the data subset from populations facing the environmental sequences with higher autocorrelation at lag 2 (sequences 25, 31, 32, 33) from both reliable and unreliable treatments for a total of 24 time series. We reserved the remaining 12 time series for cross-validation of the model fitting.

We consider two modes of transgenerational inheritance, one which is revealed by the specific parameters for combinations of offspring, maternal and grand maternal environments, which we generally call AME, for anticipatory maternal effects. Environmental effects on offspring generations can be thought of as phenotypic plasticity, while maternal and grand maternal effects have been variously called transgenerational phenotypic plasticity or deterministic transgenerational effects, see discussion in (Proulx and Teotónio, 2017). In the second model of transgenerational inheritance we consider an (unknown) environmentally-dependent property of the maternal phenotype that carries over into the next generation in a diluted fashion and that is modified by the environment where development takes place. This model is well suited to mechanisms such as inheritance of epigenetically marked DNA/chromatin with environmentally mediated germline reprogramming, as well as mechanisms where metabolites (including small RNAs) are transferred from mother to offspring and may be maintained by positive feedback loops. In both kinds of models, we include a fixed additive effect of the location of the experiment and of the blue light cue treatment.

We specified the models using STAN (Stan Development Team, 2018) and fit the models using RStan which performs Bayesian inference using a Hamiltonian Monte Carlo sampling to calculate the posterior probability of the model parameters given the observed data (R version 3.3.2, RStan version 2.15.1). We confirmed that all models reached convergence based on rhat values near 1 and that the MCMC chains were well mixed. We specified uninformative priors in STAN (see supplementary RMarkdown file for full R and Stan code). Fitted models were compared using both the generated likelihood values in the posterior parameter distribution and using a LOO-IC (Leave one out) information criterion Vehtari *et al*. (2017).

#### 3.4.1 Anticipatory maternal effects

For the AME models, we focus on models that allow for maternal and grand maternal environmental effects, indexed by the number of environment specific parameters in the model. Model 2 allows for an effect of the offspring environment (a direct environmental effect), while model 4 allows for maternal environmental effects and model 8 includes grand maternal environmental effects as well. In the supplemental materials we present a broader set of models (Table S1).

We modeled the growth rate data in each generation as a Gamma distribution where the mode and standard deviation of the distribution depended on the current and past environments experienced by the population. We chose to model and report the mode of the Gamma distribution because the mode is a more intuitive descriptor of the Gamma distribution and because the RStan function `gamma` takes rate and shape parameters that are easily calculated from the mode and standard deviation (Kruschke, 2014). In any case, the mean and variance of the gamma distribution are unambiguously defined by the mode and standard deviation. The mode of the Gamma distribution for the data with index *i* is given by

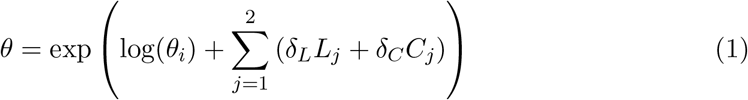

where *θ_i_* is the mode parameter specified for data point *i* for the AME model in use and given the environmental sequence (as in Table S1), *δ_L_* is the location effect and *L_i_* is the indicator variable, taking on the value of 1 for data collected in Paris and −1 for data collected in Lisbon. Similarly, *δ_C_* is the effect of the light cue given in the maternal generation and *C_i_* is the indicator variable, taking on the value of 1 for generations that received the light cue and −1 for generations that did not receive the light cue.

In more detail, we developed a set of alternative models to specify the values of *θ_i_*, that could depend on the one generation, two generation, or three generation history, and could assume asymmetric effects of anoxia and normoxia. In each model we effectively have

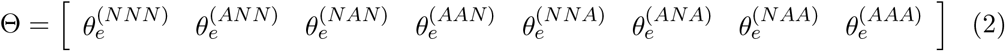

where superscripts represent the 3 generations of the environmental values leading up to and including the current environment. Thus 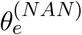 represents the modal reproductive output when the grand maternal environment was normoxia, followed by a maternal environment of anoxia, and a current offspring environment of normoxia. The values of *θ_e_* are specified in Table S1. The value of *θ_i_* is thus given by

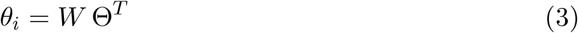

where *W* is the design vector that has values of 0 and 1 in the same order as in the definition of Θ, with 1 indicating the sequence that led up to the data point *i* and all other vector elements taking on the value 0.

The standard deviation of the Gamma distribution is assumed to depend on the offspring environment alone (*σ_A_* and *σ_N_*). Analysis of models with maternal effects on the standard deviation show little improvement (not reported).

Given the mode and standard deviation of the Gamma distribution, the standard rate and shape parameters (*α, β*) can be defined as

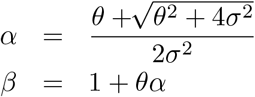

We used the function `gamma_lpdf` in *RStan* to increment the likelihood of observing each data point given the rate and shape parameters. We then used the function call `stan` run with 8 chains for 1000 iterations each, retaining the last 500 iterations for posterior probability calculations. We additionally computed LOO-IC using the package *loo* (Vehtari *et al*., 2018). To compare different models we used the `compare` function to compute the difference between LOO-IC values, divided by 2, and the estimated standard error of the difference in LOO-IC scores.

#### 3.4.2 Carryover effects

We also considered a model with a phenotype that carries over between generations. In these models there is an un-observed “epigenetic” state, *s*, that is passed on to offspring but modified by the environment every generation, *t*, which can be interpreted as germline “reprogramming” or “resetting”. The epigenetic phenotypic state simply follows a geometric decay (at a rate of *λ_N_* or *λ_A_*) towards the environment specific deviation (*c*), with the addition of a process error term independent of the environment, *ϵ*, so that in the generations experiencing normoxia the recursion is

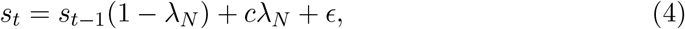

while in the generations experiencing anoxia the recursion is

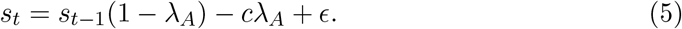

The error is modeled as a normal variate with variance 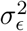 which is itself a fitted parameter in the model.

We assume a direct effect of the environment in the current generation on the population growth factor, as well as an effect of the current epigenetic state (itself carried over from the last generation and modified by current conditions). The mode for the Gamma distribution is then given by

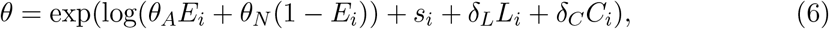

where *θ_A_* is the direct environmental effect in anoxia, *θ_N_* is the direct environmental effect in normoxia, and *E_i_* the indicator variable taking the value 1 in anoxia and 0 in normoxia. The other parameters are defined as in the AME models. Also as in the AME models, the standard deviation depends on the current environment alone and is assumed to be independent of the past environment and the epigenetic state.

### 3.5 Model validation

We performed two types of model validation, both employing posterior predictive fitting, cf. (Gelman *et al*., 2013). First, we performed posterior predictive plot analysis using the aggregate statistic of geometric mean population growth over the experimental sequences. We used this method to compare the alternative models of transgenerational inheritance to the data from which the parameters were inferred. This method allows one to identify models that miss key features of the dataset, but does not assess the sensitivity of the fit to the specific data used. We generated distribution plots of the population geometric mean fitness for AME models 1, 2, and 4 as well as for the carryover model. These were compared to the empirically observed geometric mean population growth. Our second model validation approach also used posterior predictive plots, but employed cross-validation and thus ensuring that the fit is not overly sensitive to the specific data used to fit the parameters. For cross-validation, we used the fitted models to predict time series data of the 12 populations that faced the environmental sequences 11 and 19, since these sequences had the smallest autocorrelations at lags higher than 2 when compared to the sequences employed for inference (Figures 1 and S1).

To generate posterior predictive plots, we used the `Generated Quantities` sampling capabilities of RStan to create forward simulations of the growth rates of populations facing different experimental sequences. This approach sampled parameters from the posterior distribution and then generated simulated data using the modeled probability distributions.

### 3.6 Forecasting population viability

In order to assess how the inferred transgenerational inheritance would affect population viability under different patters of future environmental fluctuations we simulated population growth under autoregressive models of environmental fluctuations. To do this we randomly drew the autocorrelation parameter, then simulated an environmental sequence of anoxia and normoxia, where the probability of repeating the same environment was *ρ*. We then filtered sequences to ensure that the number of anoxia generations was equal to the number of normoxia generations (in this way we held constant the main effect of the frequency of anoxic generations). This produced a set of environmental sequences that varied in the observed autocorrelation at lag 1 between −1 and 1, although the distribution included more negatively autocorrelated than positively autocorrelated sequences.

We then used the posterior distributions of parameters to forecast population growth dynamics, drawing a new set of parameters for each environmental sequence. We calculated the geometric mean growth rate over the entire environmental sequence.

## 4 Results

In Figure 1 we plot the observed population growth rates for the 36 populations followed during 60 generations in fluctuating environments. Table 1 shows the models’ fit and the LOO predictive density estimates, as well as comparisons between all the models analyzed. We report the elpd-LOO which can be converted to the LOO-IC score, similar to twice the log likelihood as used in model comparison using AIC (Vehtari *et al*., 2017). Because the elpd-LOO is already adjusted for the effective number of parameters in the model, values can be used for direct model comparison where larger values of elpd-LOO indicate better fitting models and the difference in elpd-LOO can be compared with the the estimated standard error of the elpd-LOO values themselves (Anderson *et al*., 1998; Spiegelhalter *et al*., 2002; Vehtari *et al*., 2017). If the change in elpd-LOO is small relative to the standard error then there is little support for the more complex model. Based on this analysis we found strong support for maternal environmental effect in our AME model, and weak support for a grand maternal effect. We also found strong support for the carryover model as compared to the AME models.

**Table 1:**
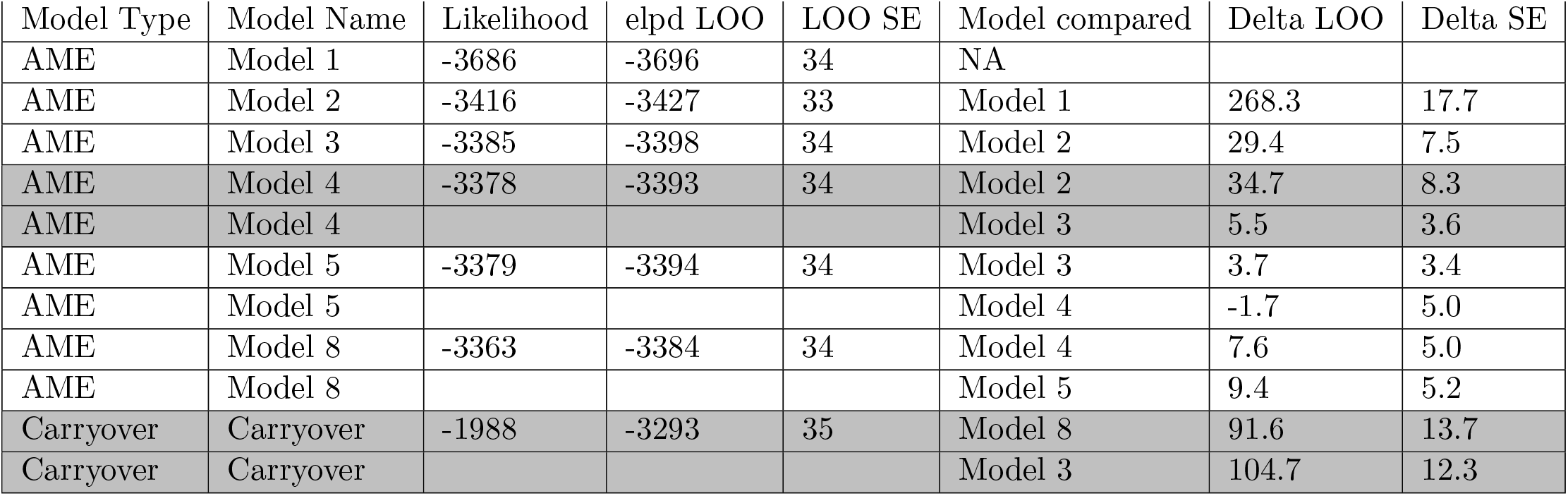
Model fit statistics. We report both the elpd-LOO and the estimated standard error of the elpd-LOO. The LOO-IC is simple-2*elpd-LOO. Model comparison implemented in the *loo* package uses elpd-LOO values that have not been multiplied by 2, and we report these directly for comparison with the estimated standard error. Highlighted models have an improvement in the elpd-LOO that substantially exceeds the standard error. AME model 4 and the carryover model were used for the additional analyses.

For all models we included an additive effect of experiment location and the blue light treatment. Estimates of these parameters were moderate and consistent between models, each having on average less than a 10% effect on the growth factor (Supplemental RMarkdown).

### 4.1 Comparisons of AME models

AME model 1 assumes no environmental effect on the mode of population growth, while model 2 assumes a direct effect of the environment in the current generation on population growth (Table S1). There is clear evidence for a direct environmental effect on population growth, with an improvement of elpd-LOO of 268.3 with a standard error of 17.7 (Table 1). Our posterior predictive plots also suggest a poor fit of model 1, with much of the observed data falling in the extreme tails of the posterior predictive distribution (Figure 4).

AME models 3 and 4 allow for effects of the environment in the maternal generation and both show a significant improvement in elpd-LOO compared with model 2 of 29.4 and 34.7 points respectively with a standard error of around 8. Model 4 also performs well in our posterior predictive plot analysis (Figure 4), with most empirically observed geometric mean growth rates falling in the region of high support. The maternal environment effect in anoxic environments are quite large (about a 50% increase), while those in normoxic environments, while distinct from zero, are smaller (about 10%, Figure 2). These results suggest that the maternal effect is stronger when the offspring experience the more stressful environment.

**Figure 2:**
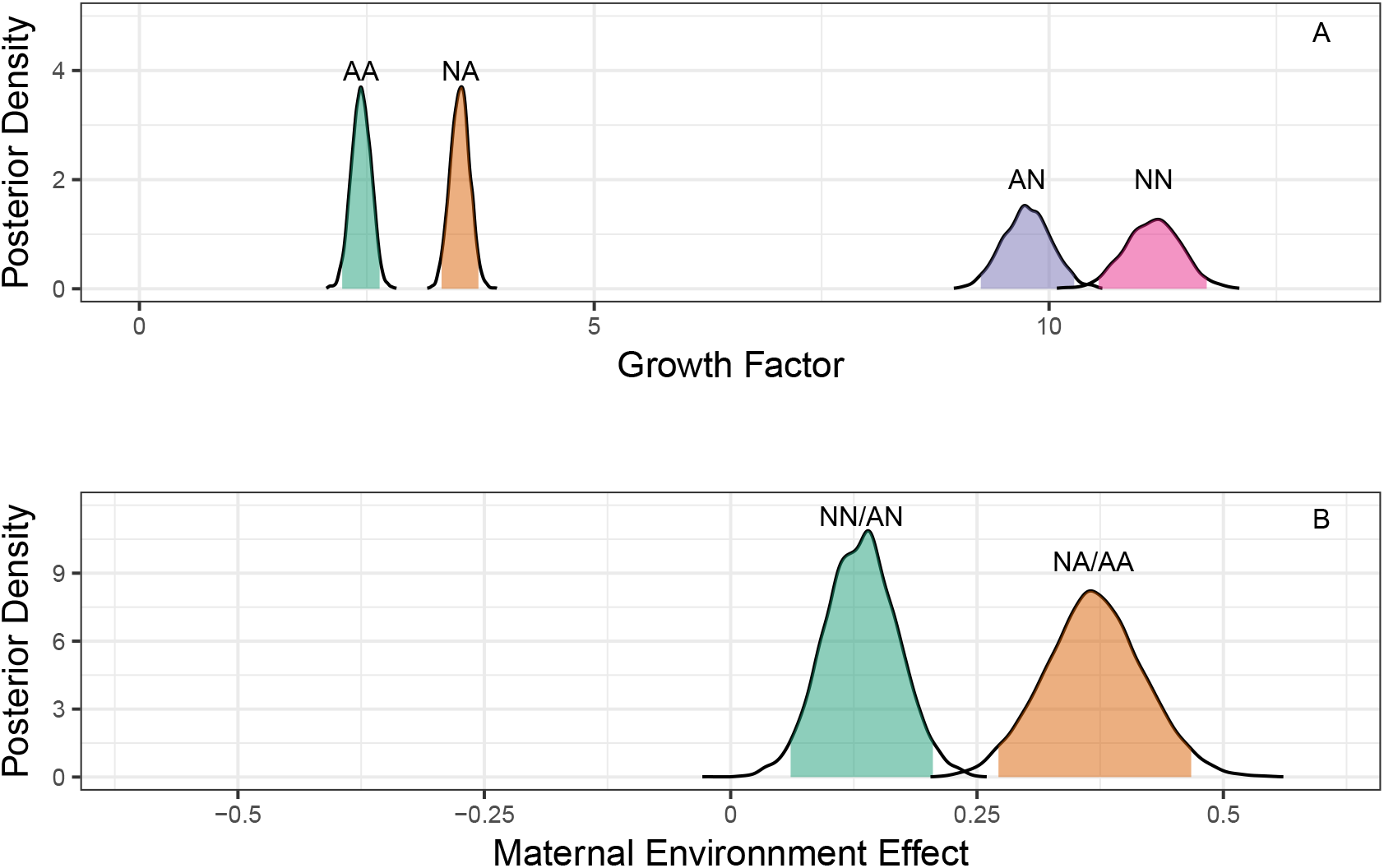
Posterior distribution information for the environment specific modes for model 4. The posterior distributions following MCMC sampling are shown, with the 95% intervals shaded. Panel A shows the environment specific effects on the mode of the Gamma distribution, which is scaled in terms of number of offspring. Panel B shows the effect of the maternal environment on the mode for individuals in anoxia and normoxia, expressed as the log of the ratio of the modes. Thus peak labeled NN/AN represents the increase These were generated by using the paired parameters samples from the posterior, thus accounting for any correlations among parameters in the posterior samples.

We further examined models accounting for 3 generation effects. Model 5 accounting for grand maternal effects when offspring face anoxia is only a marginally improvement over model 3, and not an improvement at all over model 4 (Table 1). Further inspection of the posterior parameter distributions shows that the grand maternal effect parameters in model 5 have significant overlap (Figure S4). While model 8, accounting for all combinations of 3 generations effects, does show an improvement in terms of LOO-IC over model 4 that is marginally larger than the standard error (Table 1), again inspection of the posterior parameter distributions shows that almost all grand parental parameters have strong overlap in their posterior distributions (Figure S5). One effect stands out, however, in that individuals whose grandmother experienced anoxia tend to have higher growth rates, especially when they themselves experience normoxia.

Taken together, the inference results and the model validation suggest that model 4 adequately describes the effects of transgenerational inheritance on population viability. For this reason, we use model 4 for comparisons to the carryover model below. Examination of the paired posterior distributions for model 4 (Figure S7) shows no large correlations between parameters.

### 4.2 Carryover model

Just as in the AME models, we infer a clear effect of the current environment on population growth in the carryover model (Figure 3). We also inferred a significant carryover effect in that the carryover parameter’s posterior distribution does not overlap with 0. In addition the environment-specific parameters for the degree of carryover (*λ_A_* and *λ_N_*) have a ratio that is significantly different from 0 (Figure 3C). This implies that epigenetic resetting happens more rapidly in normoxia than in anoxia, in general agreement with the AME models (Figure S8). In particular, the parameter describing the return towards the normoxic phenotypic state, *λ_N_*, is near 1, implying that there is little phenotypic “memory” in the normoxic environment.

**Figure 3:**
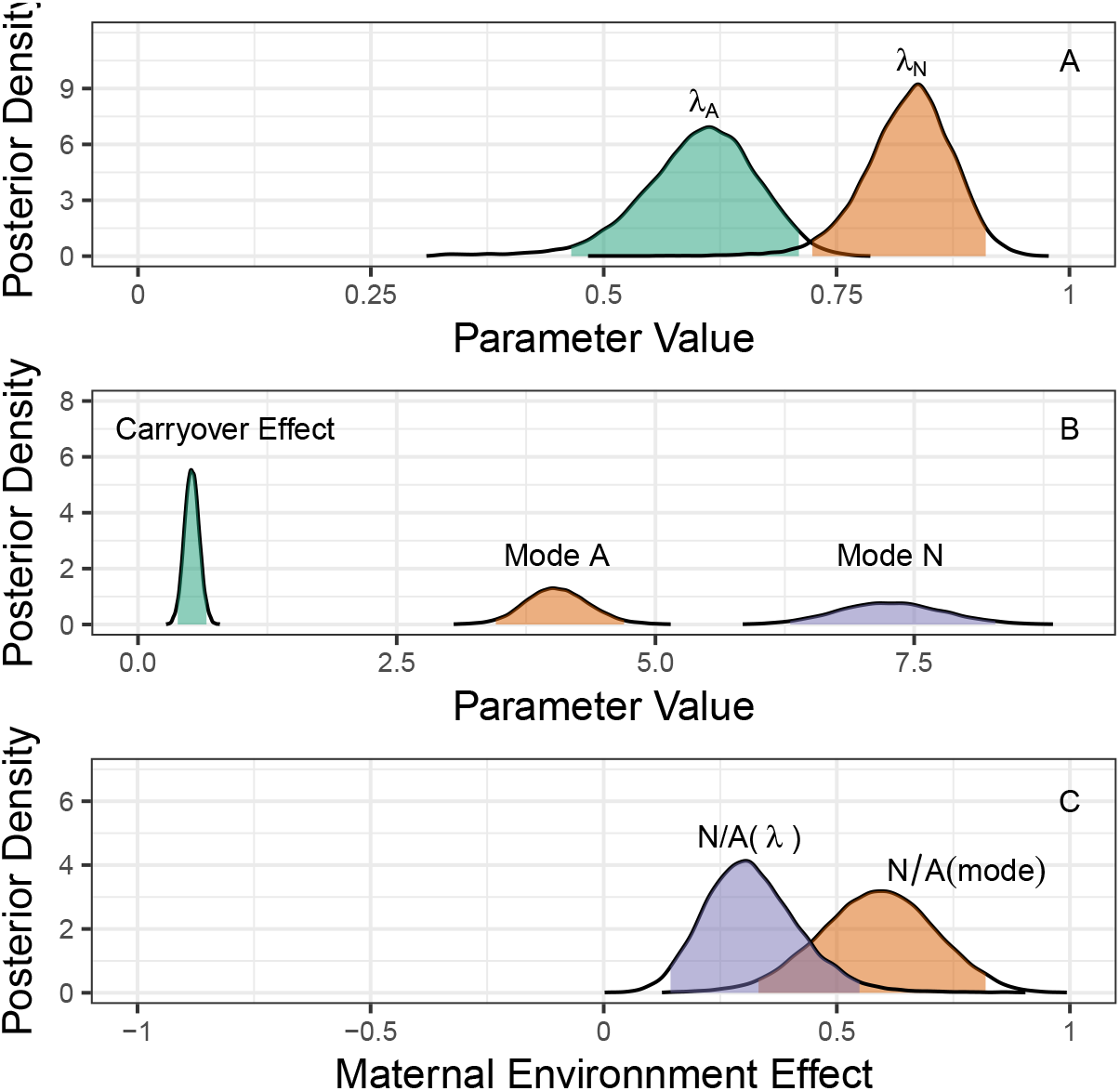
The posterior distributions for parameters from the carryover model. Panel A shows the parameters that determine the rate of carryover memory decay between generations. Panel B shows the direct environmental effect on the mode of the Gamma distribution and the magnitude of the carryover effect. Panel C quantifies the environment-specific effects. The effect of the current environment on the mode is notates as N/A(mode), while the degree that carryover depends on environment is shown as N/A(*λ*). We transform the parameters as the log ratio of the normoxic to anoxic parameter value for each posterior draw. There is a strong effect of both the current environment on growth rate and of the environment on the rate of epigenetic reprogramming.

The paired posterior distributions (Figure S9) shows that the carryover effect and the within-generation environmental effects are somewhat confounded because a larger carryover effect can compensate for lower direct environmental effects. However, our analysis of the maternal environmental effect takes these correlations into account and still indicates both significant carryover and significantly different rates of reprogramming/resetting in the two environments.

The carryover model performed well under all validation approaches, with most empirically observed geometric mean growth rates falling within the posterior distributions (Figure 4).

**Figure 4:**
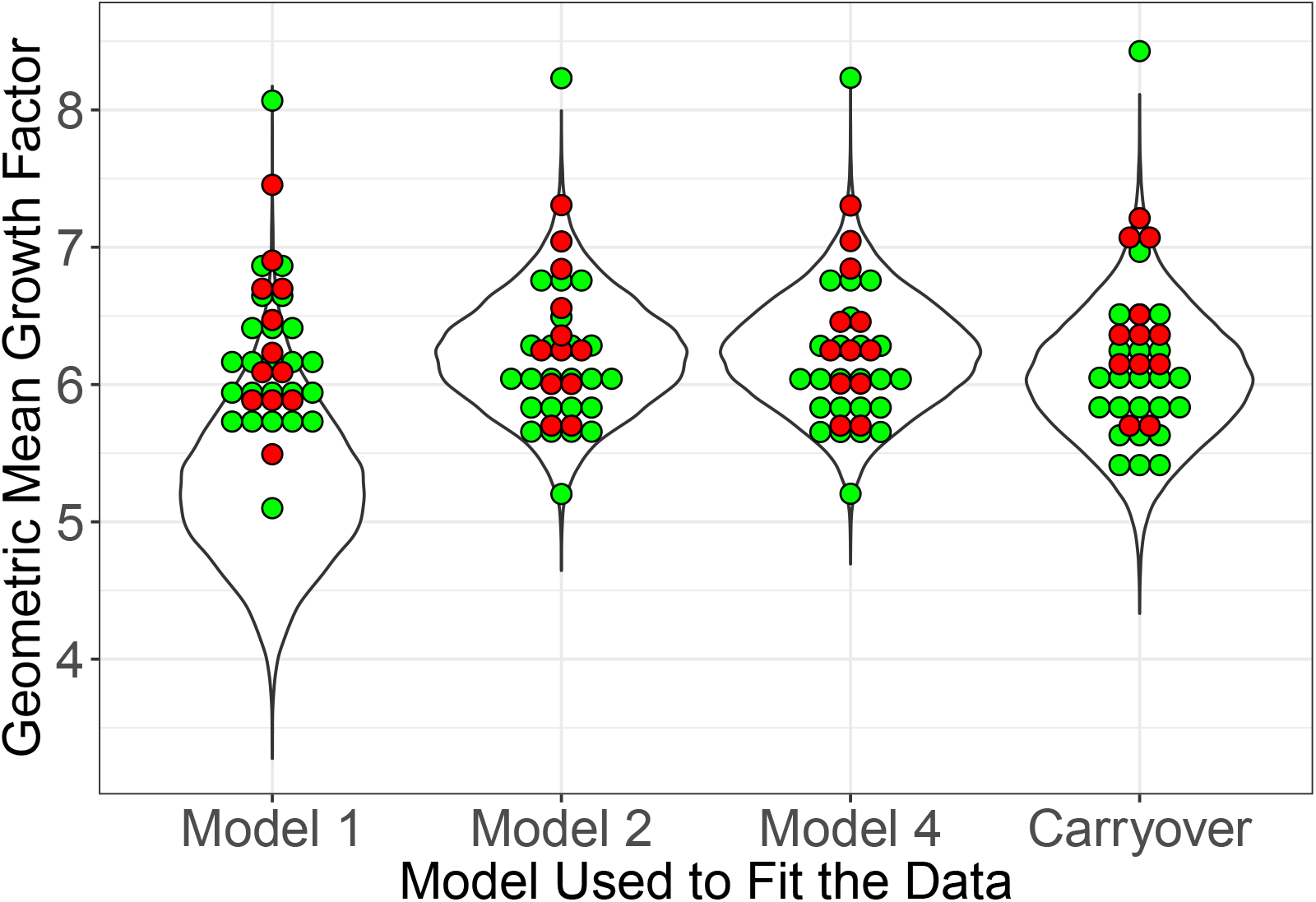
Posterior predictive plots of the different models. The violin plots show the frequency at which different geometric mean fitness values were observed. The green dots are empirically observed values from the experiments used to fit the model, with the effect of location regressed out (note that model specific location effects are used, thus the height of the empirical values differ slightly between plot elements). The red dots are the empirically observed values from experiments using sequences 11 and 19 that were not used to fit the models.

### 4.3 Comparison of AME model 4 with the carryover model

As reported in Table 1, the fit of the carryover model is a significant improvement as compared to any of the AME models. In particular, comparison of the carryover model to AME model 4 represents an improvement of 99.2 points, with a standard error of only 12.3. This is strong support for the improvement of the carryover model. We further explored the pointwise posterior prediction errors to determine which data points gave the carryover model the largest advantage over the AME models (Vehtari *et al*., 2017).

We found that the carryover model performed better than the AME models for data points that came from longer runs of a single environment, reinforcing the idea that the carryover model captures subtle effects of longer term historical effects (Figure S12).

Even though the carryover model has much higher support than the AME model 4, they produce similar outcomes in terms of their general effects. This can happen because the carryover model does a better job of predicting the variability in the data, even if both models predict similar average responses. To illustrate this, we compared the population growth rate predicted by the two models when a population experiences a repeating sequence of two generations of normoxia followed by two generations of anoxia. We sampled parameters from the posterior distribution and simulated the population growth and then summarized the data based on how many generations in a row of anoxia or normoxia had occured. The pattern of growth factor change is quite similar for the two models, and also shows a reasonable quantitative fit to the empirical data (Figure S11).

### 4.4 Forecasting population viability

We used the inferred parameters to conduct simulations of the geometric mean fitness in a range of possible environmental variability patterns. We found that regardless of the mode of inheritance, the main effect of the environmental autocorrelation at lag 1 is that increased positive autocorrelation causes a decrease in geometric mean fitness (Figure 5). This is because multiple anoxia generations in a row decreases fitness more than multiple normoxia generations in a row increases fitness. Despite the inferred carryover effects for longer than two generations, we find no relation between the geometric mean growth rate and the environmental autocorrelation at lag 2, when the environmental correlation at lag 1 is fixed at zero (Figure S13).

**Figure 5:**
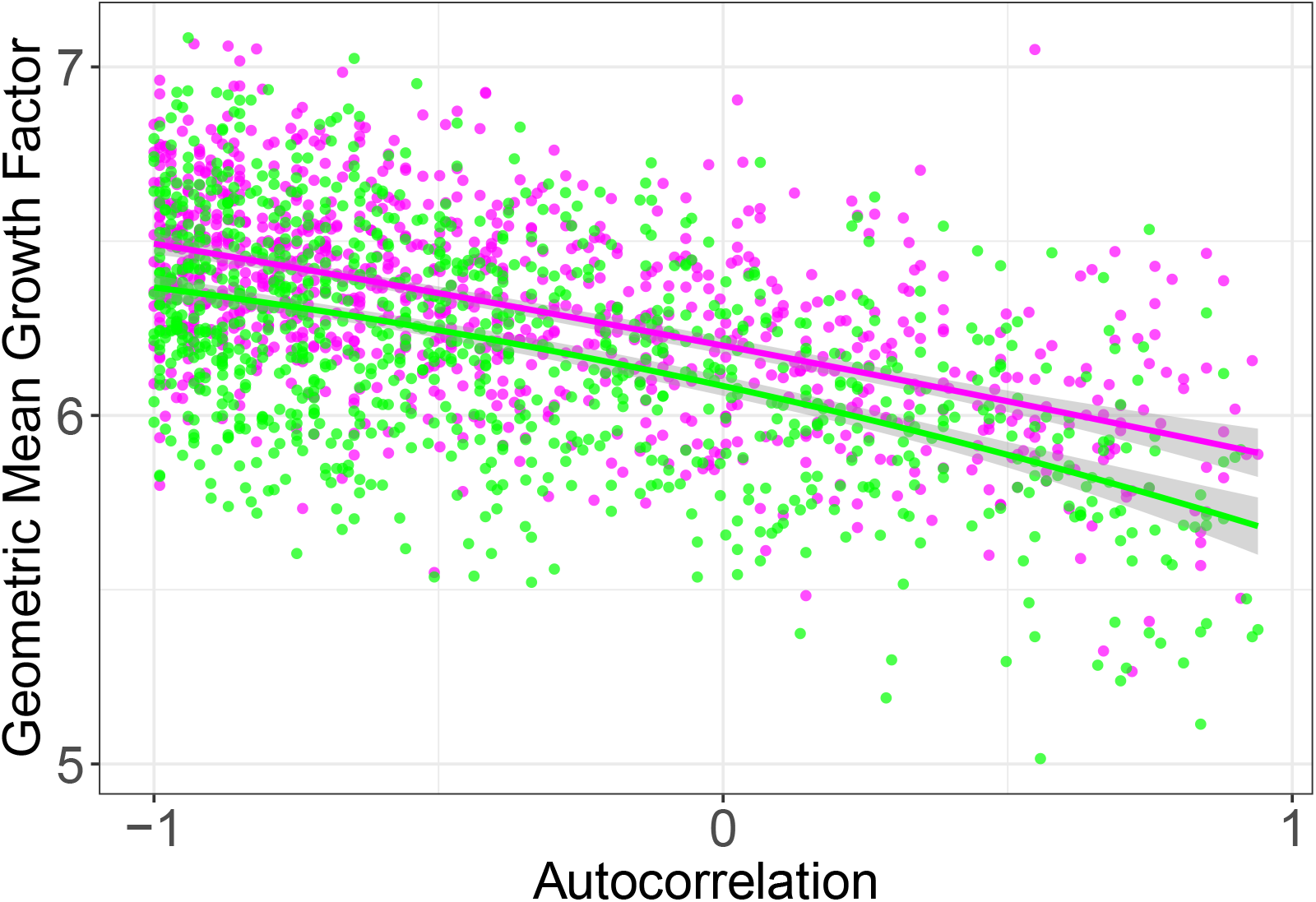
Population viability in response to the type of environmental variation, as defined by the autocorrelation at lag 1. We calculated the geometric mean growth rate from forward simulations using AME model 4 (magenta) and the carryover model (green). Curves are quadratic fits, with shaded areas around them the 95% credible intervals.

## 5 Discussion

We employed a highly replicated set of experiments to fit competing models of trans-generational inheritance in order to explain population growth dynamics in fluctuating environments. In the experimental life cycle, larval to adult population size was held constant while populations were exposed to fluctuating oxygen level conditions during embryogenesis and initial larval growth. This design feature allowed us to separate the effects of the environments where development until maturity takes place from those of transgenerational inheritance of maternal phenotypes.

Similar approaches to ours have long been used in natural and experimental populations to infer the effects of population subdivision (including age and life history stage structure), competitive exclusion, predation, pathogen infection or resource allocation, e.g., (Mueller and Joshi, 2000; Lande *et al*., 2003, 2006; Chevin *et al*., 2015). Hence, one venue for future research is to understand if transgenerational inheritance could explain population dynamics in tandem with within-generation effects (Ruokolainen *et al*., 2009; Inchausti and Ginzburg, 2009). In natural populations, anticipatory maternal effects are known to affect population and species community dynamics, e.g., (Dantzer *et al*., 2013; Duckworth *et al*., 2015), but is usually the case that maternal density itself is the cue for the expression of transgenerational inheritance, making it difficult to disentangle the several sources of density regulation. Experimental models such as ours offer the possibility of manipulating population sizes from initial larval growth to adult reproduction (which here was held constant), together with the environmental conditions during embryogenesis and initial larval growth that induce transgenerational inheritance.

Our analysis provides strong evidence for an effect of the maternal and the grand maternal environment. First, AME models that include an influence of the maternal environment have much better likelihood and LOO-IC scores. Second, our AME models with maternal environmental effects shows distinct posterior distributions for the inferred fitness affects in the four possible pairs of maternal-offspring environment. For offspring facing either environment, population growth is higher when the maternal environment is the permissive normoxic environment. Third, we found even stronger support for our carryover model that allows for multi-generation accumulation of the environmental history. Each of these points towards the presence of transgenerational inheritance of phenotypes that persist at least through the maternal and grand maternal generations. Differentiating anticipatory maternal and grand maternal developmental and physiological mechanisms as defined in AME models from carryover mechanisms is an empirical challenge, and is difficult because AME and carryover will only differ in the degree of phenotypic expression (penetrance) at the individual level of analysis and in the fraction of the progeny that will inherit maternal phenotypes at the population level of analysis.

In *C. elegans*, anoxia greatly reduces survival if maternal generations are cultured in a hyperosmotic environment and hermaphrodites cannot provision their embryos with sufficient glycogen (Frazier and Roth, 2009; Dey *et al*., 2016). It is therefore not surprising that AME models, particularly model 4, provides a good fit to the observed population growth dynamics. We are, however, currently ignorant about the transgenerational mechanisms that could explain the good fit of the carryover parameters. It is known that anoxia exposure during embryogenesis results in cell-cycle arrest and chromosome reallocation to the nuclear lamina, which in turn leads to chromatin changes and to reductions in gene expression, all presumed to be energy saving physiological responses enhancing survival (Padilla *et al*., 2014). It is unknown if some or all of these molecular and cellular responses to anoxia are inherited between generations, and if so in a quantitative way compatible with the carryover model. As with other environmental challenges, such as starvation and high temperature (Houri-Zeevi and Rechavi, 2017; Minkina and Hunter, 2018), one can speculate that small RNA gene silencing and/or histone methylation modifications are inherited for several generations after an anoxia episode.

We further found evidence for an asymmetric effect of history in the stressful versus the more permissive environment. For AME model 4, the effect of the maternal environment when offspring are in anoxia is about twice as large as that in normoxia (Figure 2). The carryover model further distinguishes these effects, showing faster germline reprogramming in normoxia than in anoxia (Figure 3). Based on the mean values of the posterior distributions for the *λ* values, it takes about 3 generations for complete resetting in normoxia, but about 6 generations for complete resetting in anoxia (Figure S8, as measured by achieving 99% of the asymptotic value). Little empirical data supports this asymmetry in germline reprogramming in *C. elegans*. One exception is the study of Frézal *et al*. (2018), where genotypes prone to progressive sterility with more and more generations at a high 25°C temperature (a temperature that is physiologically stressful since broods are small), can recover fertility and be cultured indefinitely when exposed for a single generation to the permissive 20°C, a temperature where hermaphrodites have regular brood sizes. Asymmetric germline reprogramming is perhaps a common feature of all organisms in that keeping a memory of past challenging environments may allow rapid resource allocation for survival when those environments are predictably encountered in the near future, while near-complete reprogramming and resetting may allow rapid resource allocation for reproduction as soon as environmental conditions become favorable.

Our comparison between AME and carryover models involved comparing models that are not nested. To understand why the carryover model produced such higher likelihood and LOO-IC scores we also examined the point wise LOO values. These represent the contribution of each data point to the overall LOO score and can be used to develop appropriate models for the data (Vehtari *et al*., 2017). We found that the carryover model consistently performs better in predicting population growth during generations where the history has included multiple generations of the same environment (Figure S12). This suggests that the AME models do not capture longer term history, and in fact may suffer from idiosyncratic over-fitting due to unplanned correlations within the environmental sequences that we used. On the other hand, the carryover model is responsive to variation in the environmental history using only a few parameters, and therefore is less at risk of over fitting.

Even though the carryover model provides a better fit to the data, the fitted AME and carryover models produce similar predictions, both in terms of environment specific fitness (Figure S11) and in terms of forecasting population viability (Figure 5). Our forecasting of population viability depends on measuring the density-independent growth rates. In nature, when populations are already at risk of local extinction we do not expect significant effects of density-dependence, and further expect that any density-dependent feedback would simply make populations more vulnerable. Using the fitted models, we found that populations are expected to show reduced viability in environments that are positively autocorrelated, when the mean frequency of the two environments is held constant. This effect is due to the increase in runs of generations in the same environment. Runs of anoxic generations have a larger negative effect on population growth than the positive effect of runs of normoxic generations. This result is likely general, in that it only depends on the more extreme response to stressful environments. While the effects of transgenerational inheritance on population viability are modest, on the order of 15%, this magnitude of difference could be important for at-risk species.

Long-term population viability is mostly a function of the specific sequence of environmental fluctuations that that particular population will face in the near future (Figures 5 and S13). However, AME model 4 consistently predicts on average about 3% higher population viability than the carryover model, regardless of the degree of autocorrelation in the environment. Thus for the specific parameters inferred here, we expect stronger selection for genotypes expressing anticipatory maternal effects than for genotypes expressing carryover effects. Even if selection for strictly (within-generation) phenotypic plasticity (Figures 2 and 3) is expected to be much stronger than for transgenerational effects, cf. (Kuijper and Hoyle, 2015; Proulx and Teotónio, 2017). The predicted selective differences between differing kinds of transgenerational inheritance could determine the adaptive potential of populations in fluctuating environments and the probability of evolutionary rescue when at the brink of extinction (Martin *et al*., 2013; Chevin *et al*., 2013).

In conclusion, we have found that differing modes of transgenerational inheritance can affect population viability in fluctuating environments, and more specifically that an yet undescribed carryover mechanism in part determines population growth rates in fluctuating oxygen level conditions in *C. elegans*. Asymmetric germline reprogramming, in that the most challenging anoxic environmental effects persists for longer than the benign normoxic effects, explains why population viability is negatively related to the degree of environmental autocorrelation. Further study will be needed to understand the relative effects of within and between generation factors in population density regulation.

## 6 Acknowledgments

We thank T. Vale for building the LED plate-racks and writing LUCS, P. Sandner for preliminary light sensitivity assays, I.M. Chelo, P. Ibañez, H. Gendrot, J. Nedli, S. Nunes and S. Santos for help with worm handling, and L.-M. Chevin, V. Colot, M.-A. Félix, L. Frézal, G. Lepère and L. Noble for discussion. Data analysis was done at the Center for Scientific Computing at UC Santa Barbara, an National Science Foundation (NSF) supported facility (DMR-1121053, CNS-0960316). SD was a fellow of the Labex MemoLife program (ANR-10-LBX-54 MEMO LIFE and ANR-IDEX-0001-02-PSL). The project was funded by grants from the NSF (EF-1137835) to SRP, and the European Research Council (FP7/2007-2013/243285) and Agence Nationale de la Recherche (ANR-14-ACHN-0032-01) to HT.

## A Supplementary Figures

**Figure S1:**
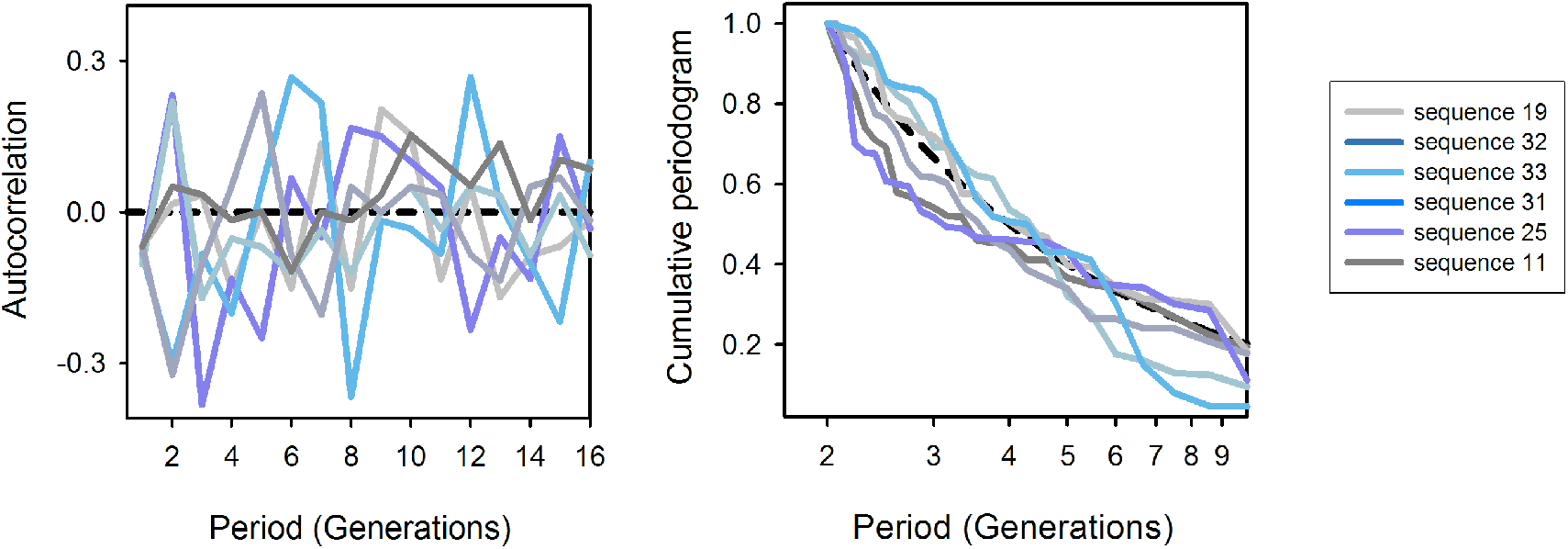
Left plot shows the autocorrelation function of the 6 environmental sequences of normoxia and anoxia oxygen level conditions employed in the experiments, as calculated with the acf function in R. We designed sequences controlling for the probability of environmental conditions being repeated between maternal and offspring generations (*ρ*_1_) and between grand maternal and offspring generations (*ρ*_2_), each scenario being represented twice, with 6 replicate populations facing each one of them (total of 36 populations followed during 60 generations). Right plot shows the Fourier transform of the autocorrelation functions of the environmental sequences, as the cumulative sine periodogram over period. We used the spectrum and cpgram functions in R to calculate these periodograms (Venables and Ripley, 2002). The pattern of environmental fluctuations can be characterized in terms of the light spectrum decomposition, with predominance of short period fluctuations indicating “blue” environments and predominance of long term fluctuations indicating “red” environments (Ruokolainen *et al*., 2009). Lack of a predominance of fluctuations over any period is shown as a dashed line.

**Figure S2:**
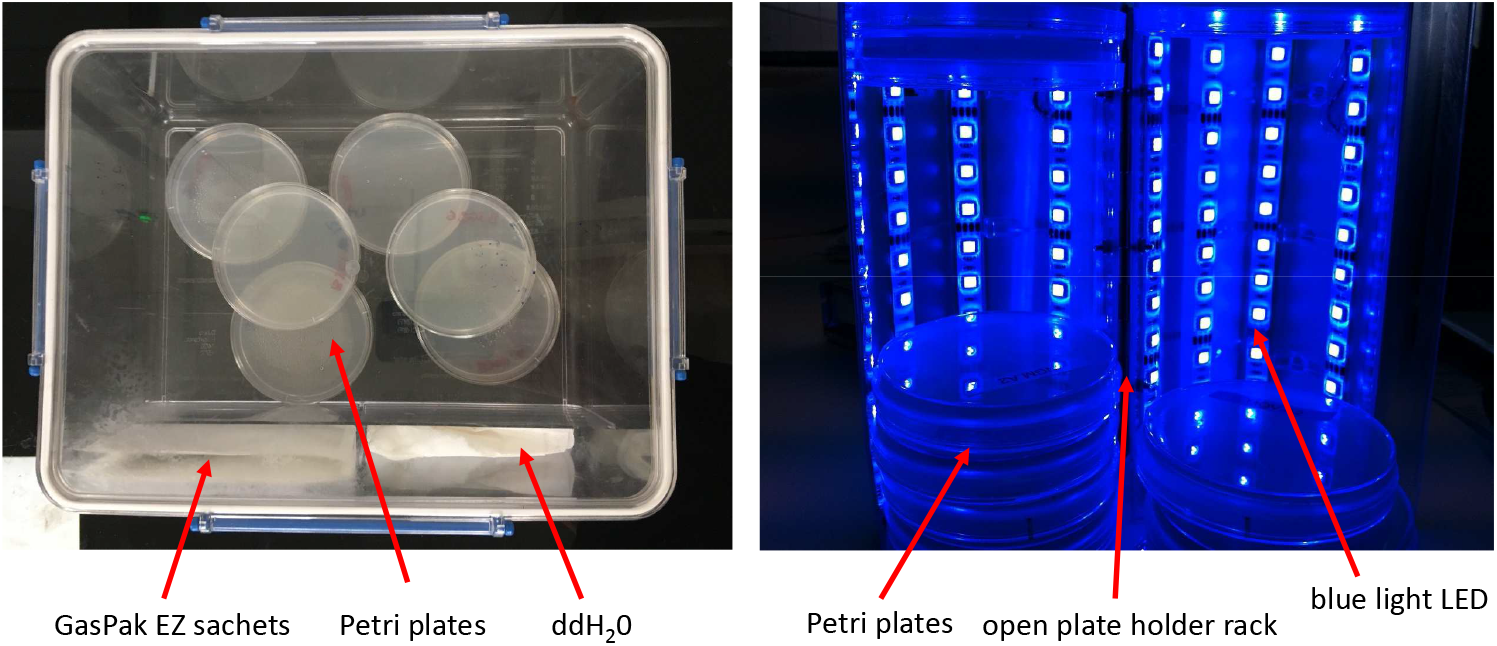
Photographs of the apparatus employed in the experiments. Left, boxes containing Petri dish plates with embryos obtained from harvesting 10^4^ adult hermaphrodites from each population, without *E. coli* food. Employment of GasPack EZ sachets ensured anoxia conditions during embryogenesis and larval growth until starvation-arrested L1. After 16h-18h in anoxia, or in normoxia conditions, 10^3^ live L1s were seeded into each of 10 Petri dish plates with food, per population, for initiation of a new generation. For three days these plates were maintained under our regular lab protocols, except that starting 48h after L1 seeding, and for a period of 12h, maternal hermaphrodites were exposed to a reliable or unreliable blue light cue of offspring anoxia conditions (right panel).

**Figure S3:**
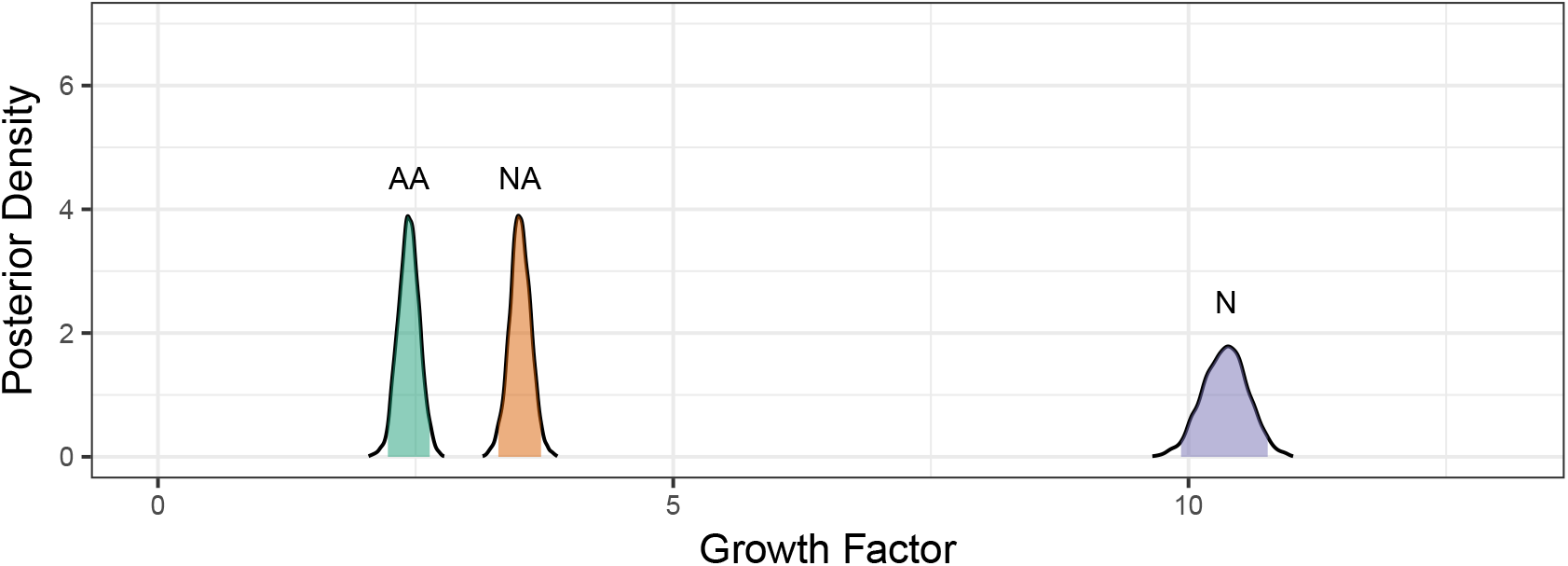
The environment specific modes for population growth rate for AME model 3. The posterior distributions following MCMC sampling are shown, with the 95% intervals shaded.

**Figure S4:**
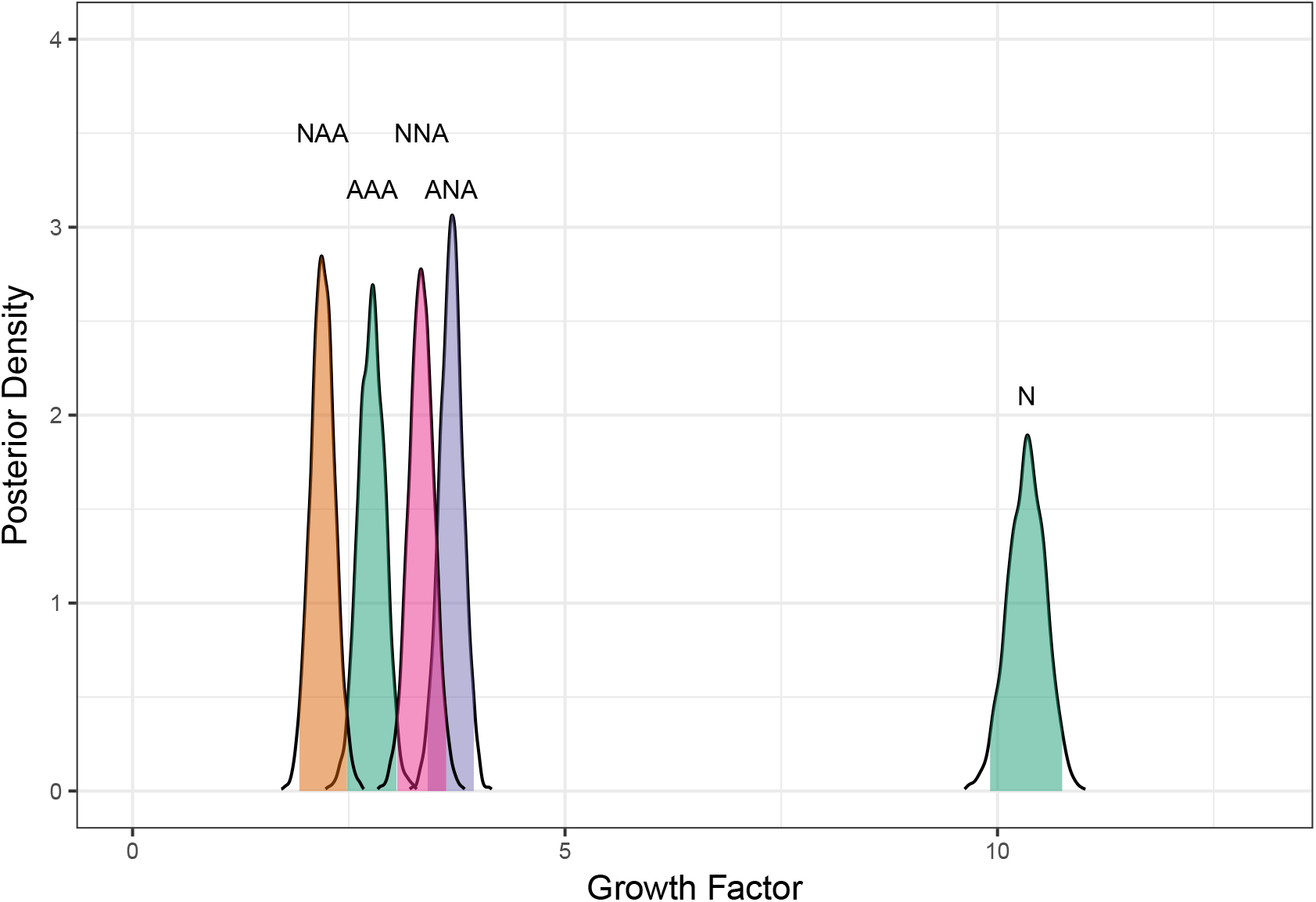
The environment specific modes for population growth rate for AME model 5. The posterior distributions following MCMC sampling are shown, with the 95% intervals shaded.

**Figure S5:**
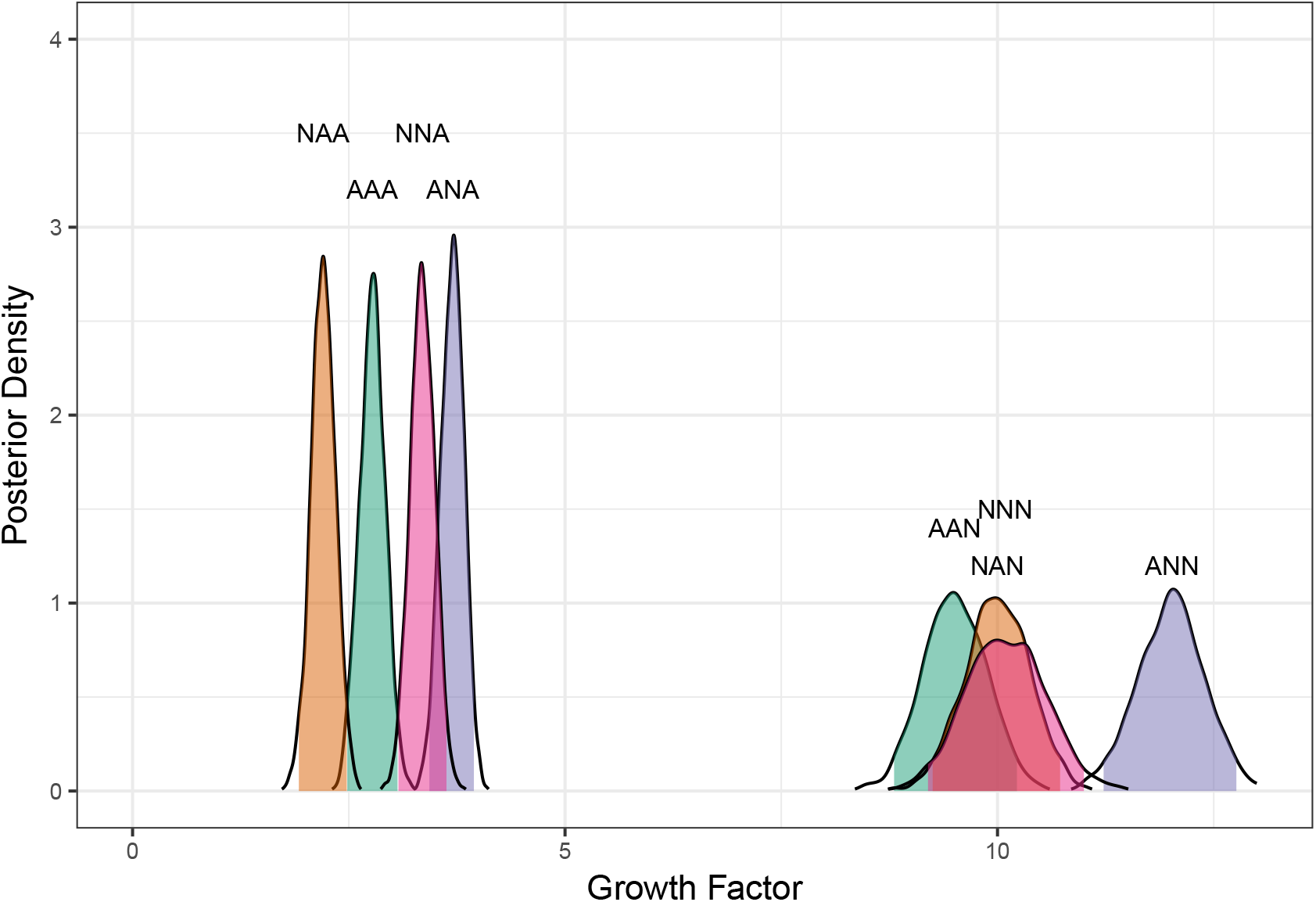
The environment specific modes for population growth rate for model 8. The posterior distributions following MCMC sampling are shown, with the 95% intervals shaded.

**Figure S6:**
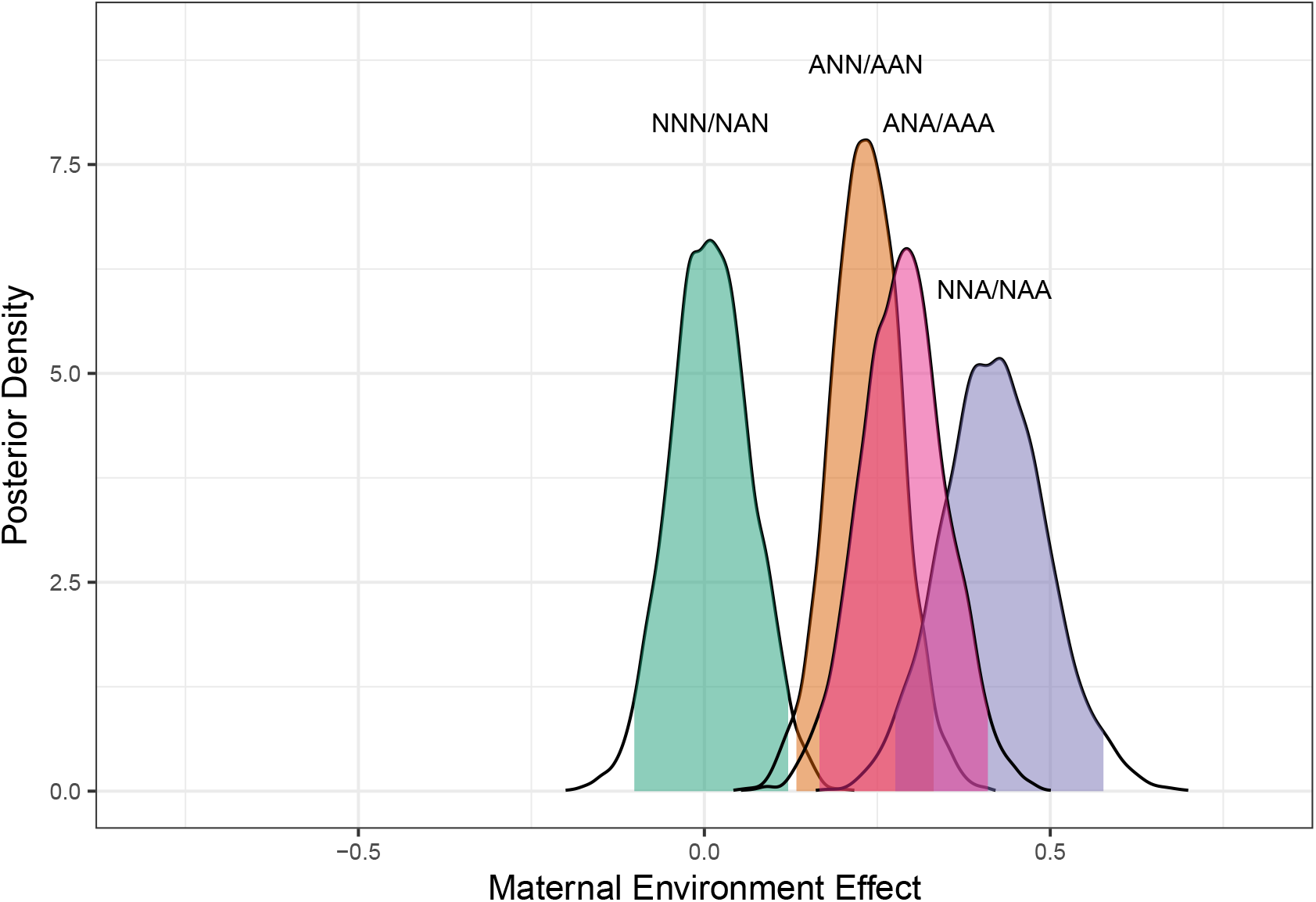
Model 8 effects of maternal environment when offspring are faced with anoxia and normoxia. We took the log of the ratio of the mode of reproductive output when the parents faced normoxia to anoxia for the posterior distribution samples and plotted their densities. Peaks are labeled by the contrasted modes.

**Figure S7:**
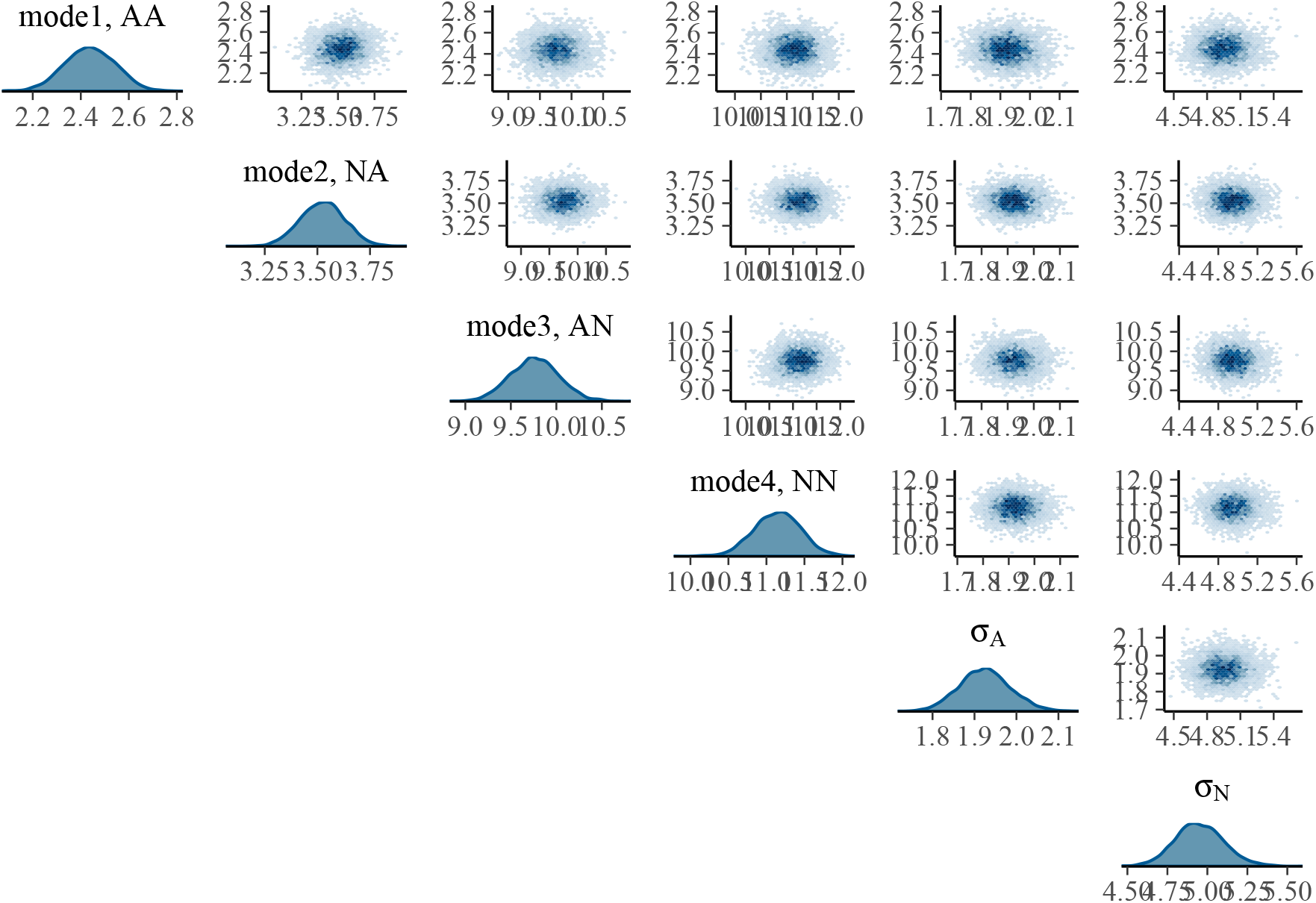
AME model 4 joint posterior parameter distributions are shown in the off diagonal plots. The diagonal plots show the parameters as in main Figure 2, together with offspring environment standard deviations, *σ_A_* and *σ_N_* (note the much reduced growth factor variance in anoxia when compared to normoxia).

**Figure S8:**
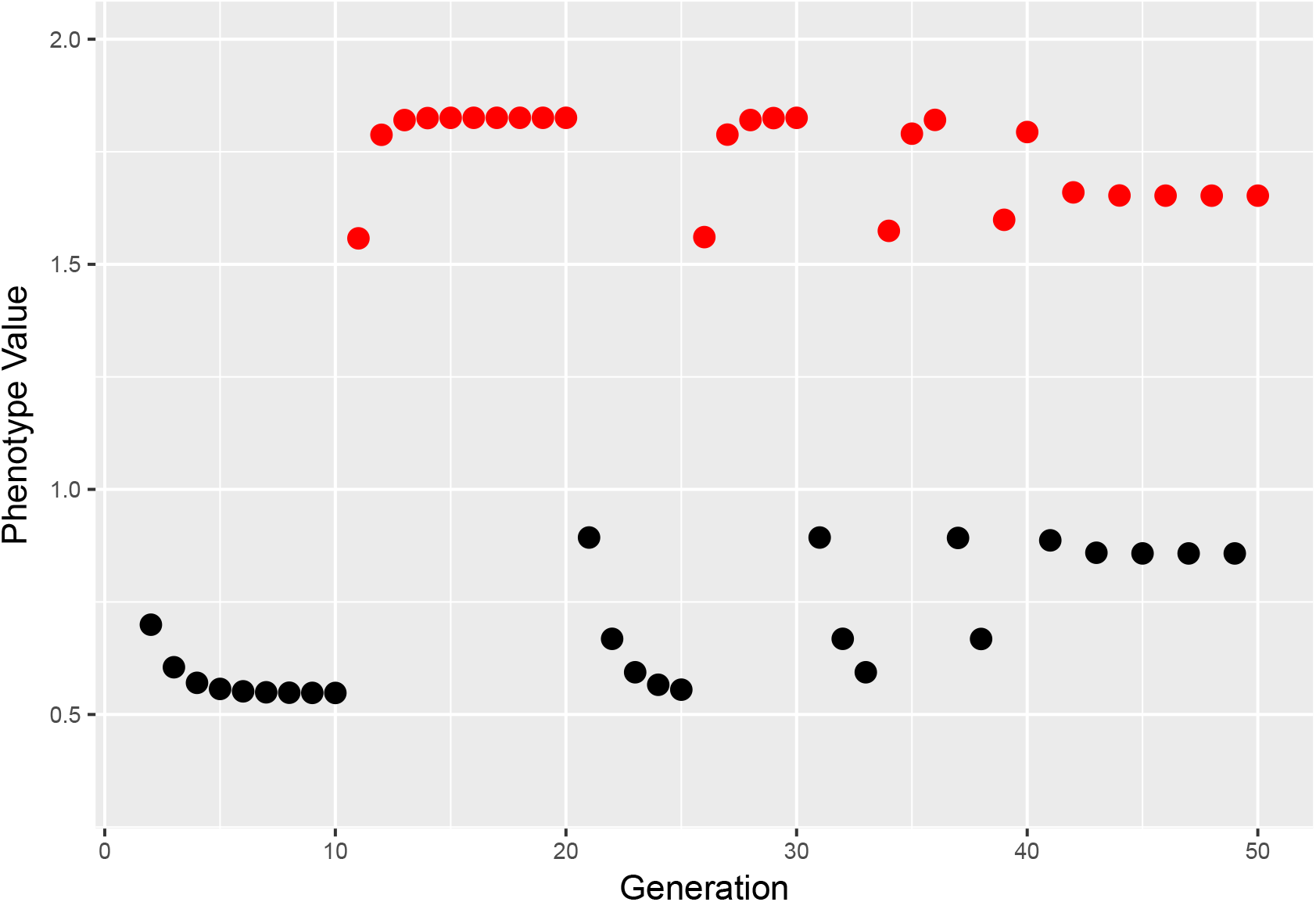
Illustrative plot of the carryover effect using the mean parameters from the posterior distribution of the model fit. The imposed environment is indicated by the color coding where black is anoxia and red is normoxia. A switch from normoxia to anoxia shows an effect of the environmental history for several generations, while a shift from anoxia to normoxia shows an effect of the maternal environment, but little effect of prior generations.

**Figure S9:**
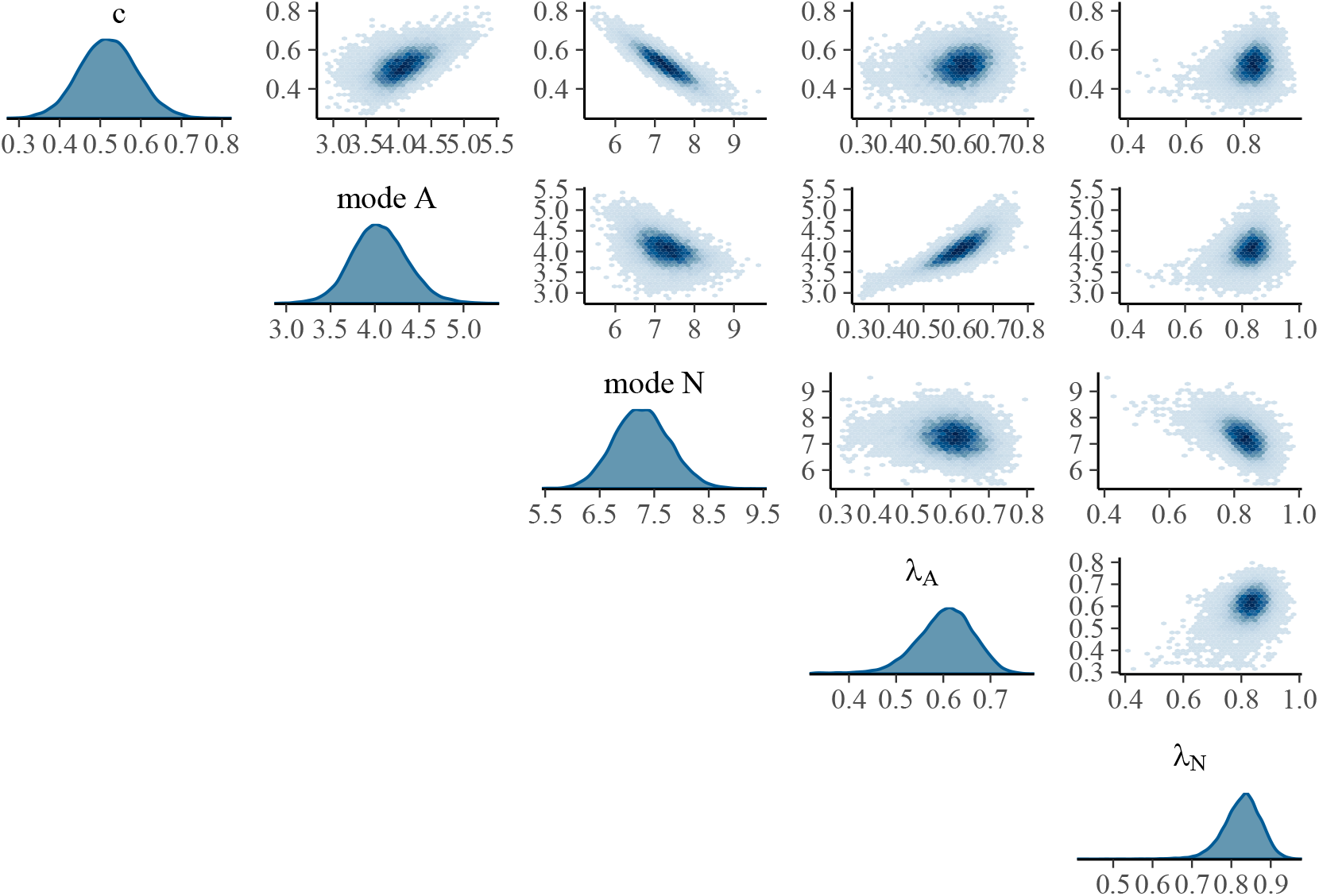
Carryover joint posterior parameter distributions are shown in the off diagonal plots. The diagonal plots show the posterior distributions as in main Figure 3.

**Figure S10:**
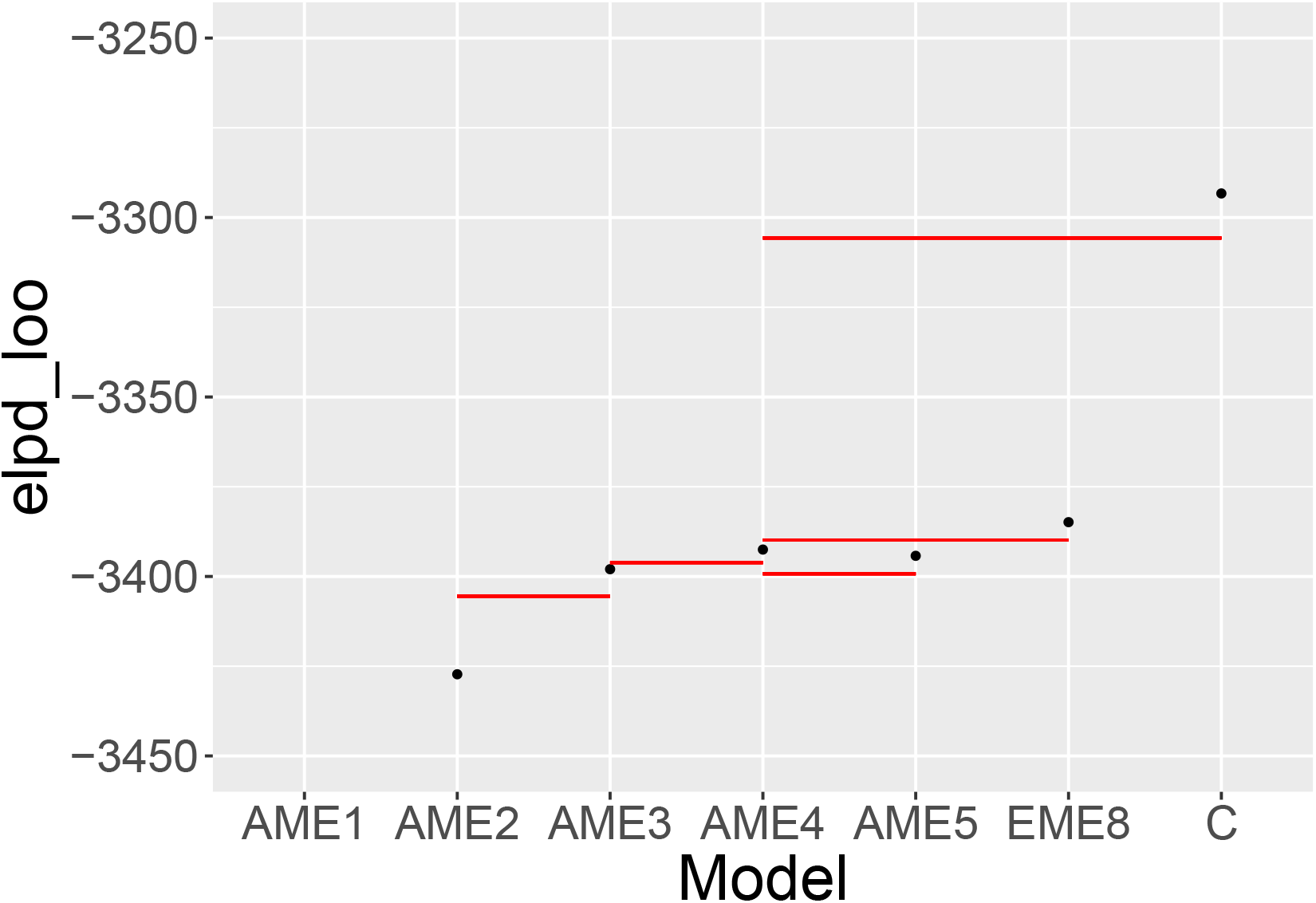
The elpd-loo scores for the different models. The dots represent the elpd-loo estimates for each model, while the red lines represent the standard error of the difference in the LOO estimates between compared models. Because this value depends on both models being compared, it is shown to highlight the most important comparisons. The line is drawn from the more complext model being compared towards the left to indicate which model is being compared. If the red line is above the point on the left side of the line, then there is an advantage to the more complicated model.

**Figure S11:**
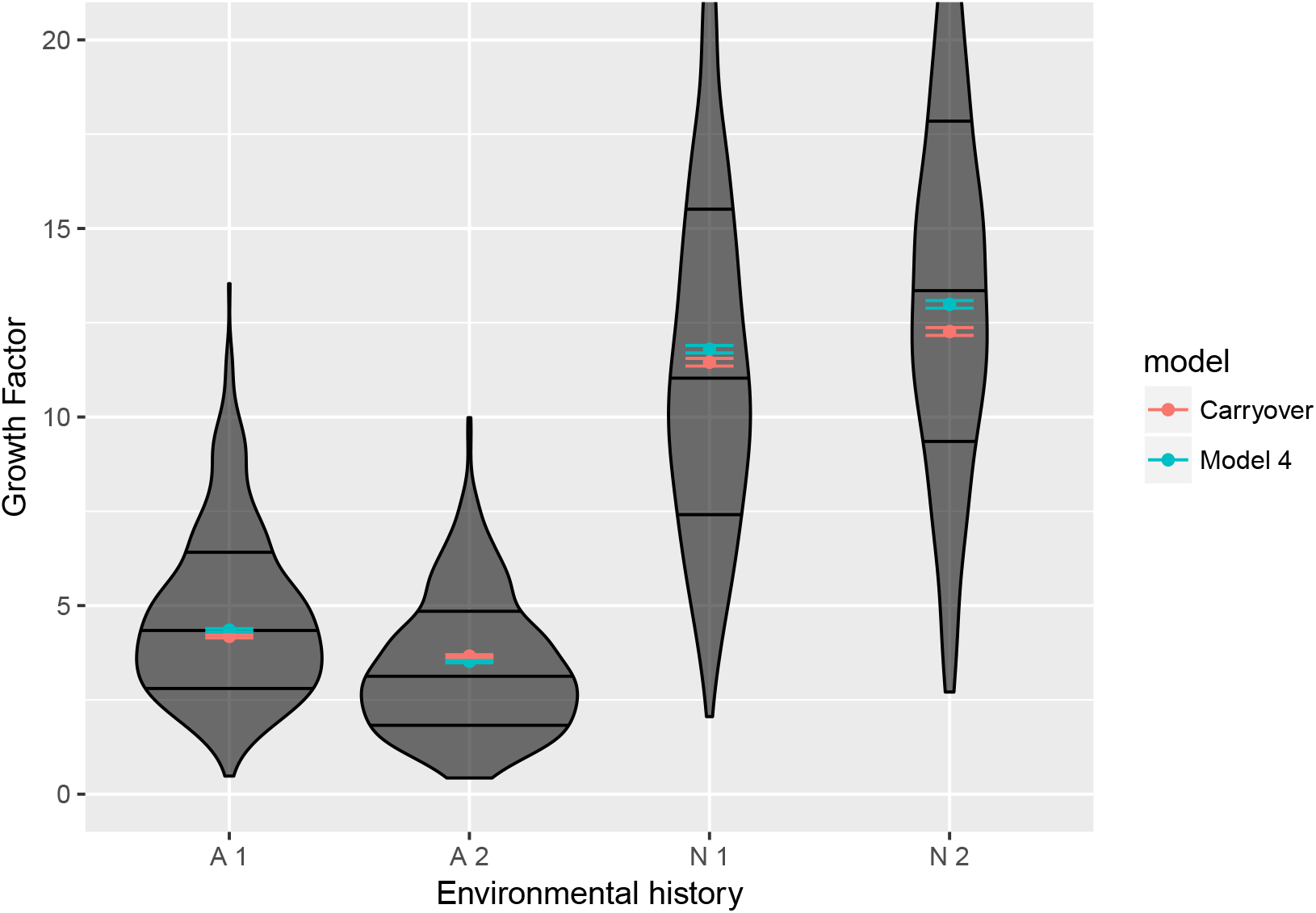
The environment specific growth rate outcomes of the AME and carryover models. We considered a repeating sequence of N-N-A-A environments and sampled parameters 500 times from the posterior distribution. We calculated the growth factor after one or two generations in Anoxia and Normoxia (A1, A2, N1, N2). Overlaid are the empirically observed distributions of the growth factors, where we have grouped all observations based on whether it is the first generation in that environment. Thus the empirical data have heterogeneity in the history earlier than the maternal generation which is not present in the simulated data. The location values were regressed out of the empirical data.

**Figure S12:**
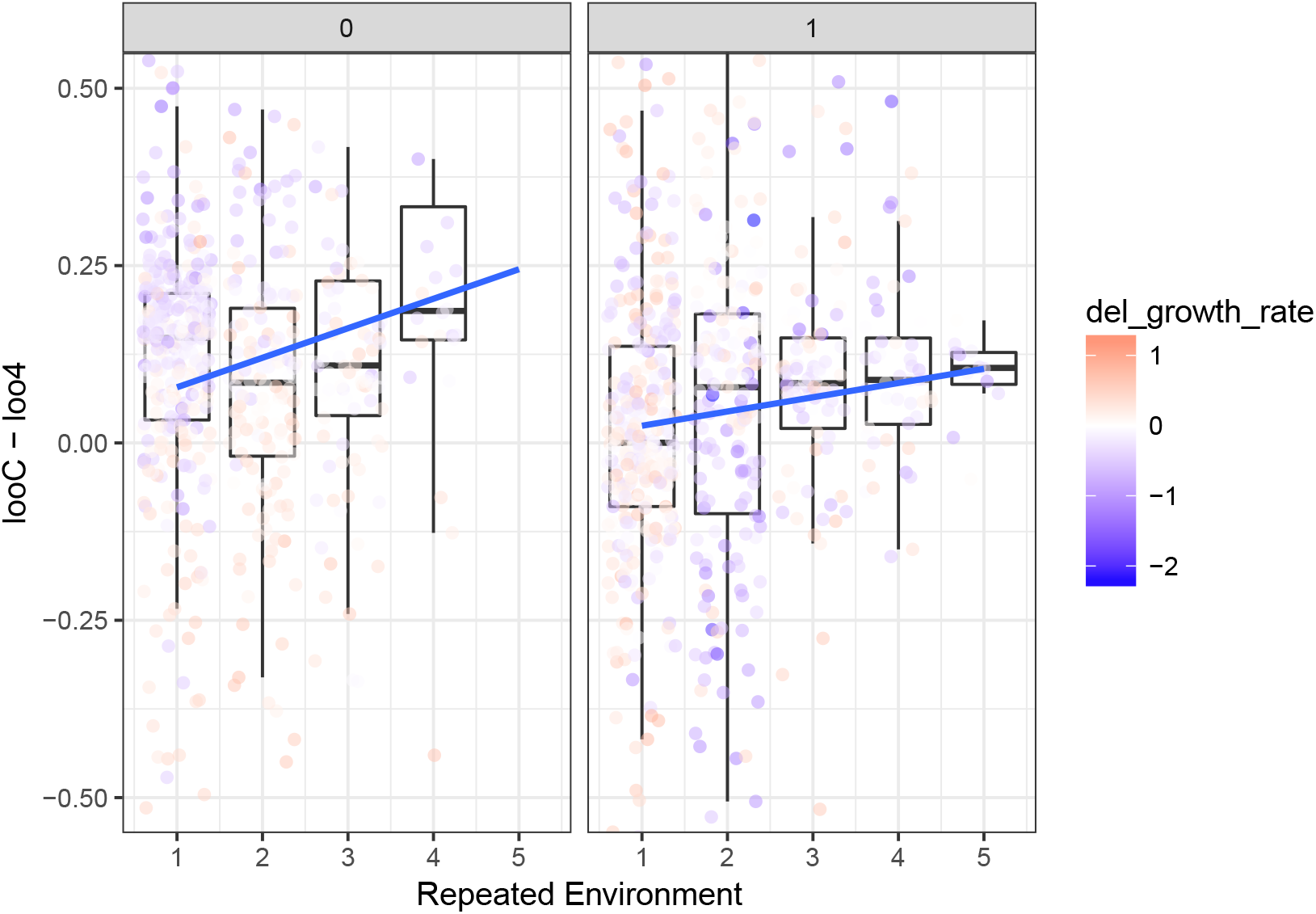
Plots of the difference between pointwise LOO values for the carryover model and AME model 4. We grouped data by the number of generations in a row that preceded the focal generation and extracted the pointwise LOO density. The plots show the difference between the pointwise LOO density for the carryover model and model. This difference in pointwise LOO values between the models represents the degree to which the fit of a single point contributes to the LOO score of that model. Larger positive values mean that the point contributes more to the total difference between the carryover model and model 4, in effect signaling that the fits of these points are the points most responsible for the overall advantage enjoyed by the carryover model. Data are separated by the current environment, where 0 is normoxia and 1 is anoxia. Points are colored to represent the degree to which the growth rate in the focal generation differed from the average growth rate in that environment. Regression lines show that increased time in a single environment produces a better fit by the carryover model.

**Figure S13:**
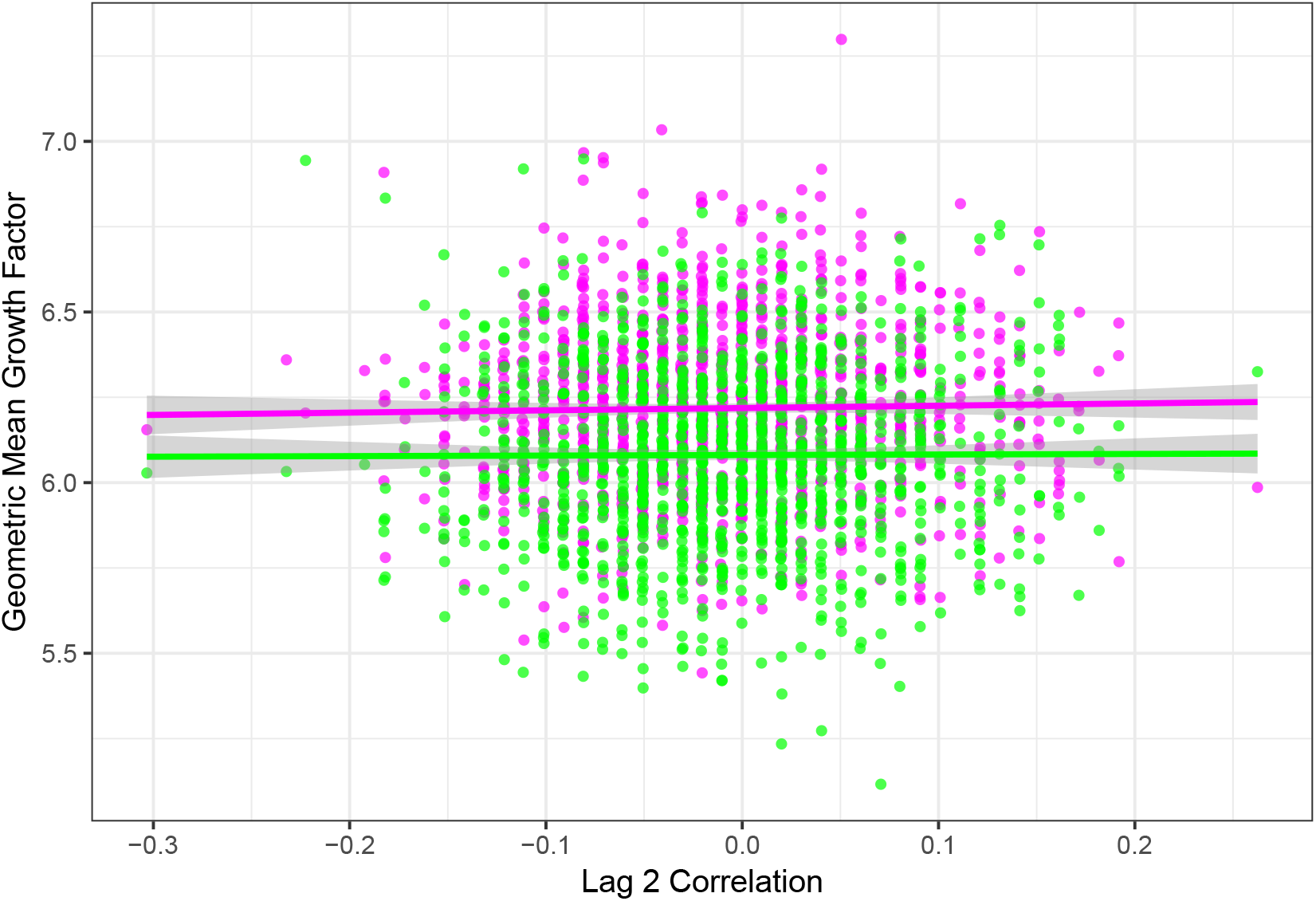
Population viability in response to the type of environmental variation, as defined by the autocorrelation at lag 2 and only for sequences where lag 1 autocorrelation is zero. We calculated the geometric mean growth rate from forward simulations using AME model 4 (magenta) and the carryover model (green). Curves are quadratic fits, with shaded areas being the 95% credible intervals.

### B Supplementary Table

**Table S1:** The table shows how the *θ_e_* parameters are related to each other. Models are named by the number of distinct parameters. Model 1 has a single parameter, *mode 1* that is the mode of the Gamma distribution for offspring production for each environmental history. Model 2 involves one parameter for offspring in anoxia and another for offsrping in normoxia. Model three allows for a parental effect when offspring are in anoxia, but not in normoxia. Model 4 includes parental effects in both offspring environments. Like Model 3, Model 5 allows for history to effect reproduction in anoxia, but not normoxia. Finally, Model 8 allows for distinct effects in each of the eight sequences of three generation environmental history. Models 6 and 7 have not been explored since they only assume grand maternal environmental effects.

| | Description | $\theta_e^{(AAA)}$ | $\theta_e^{(NAA)}$ | $\theta_e^{(ANA)}$ | $\theta_e^{(NNA)}$ | $\theta_e^{(AAN)}$ | $\theta_e^{(NAN)}$ | $\theta_e^{(ANN)}$ | $\theta_e^{(NNN)}$ |
| --- | --- | --- | --- | --- | --- | --- | --- | --- | --- |
| Model 1 | Baseline growth factor | mode 1 | mode 1 | mode 1 | mode 1 | mode 1 | mode 1 | mode 1 | mode 1 |
| Model 2 | Phenotypic plasticity | mode 1 | mode 1 | mode 1 | mode 1 | mode 2 | mode 2 | mode 2 | mode 2 |
| Model 3 | Anoxia maternal effect | mode 1 | mode 1 | mode 2 | mode 2 | mode 3 | mode 3 | mode 3 | mode 3 |
| Model 4 | Normoxia and anoxia maternal effect | mode 1 | mode 1 | mode 2 | mode 2 | mode 3 | mode 3 | mode 4 | mode 4 |
| Model 5 | Anoxia grand maternal effect | mode 1 | mode 2 | mode 3 | mode 4 | mode 5 | mode 5 | mode 5 | mode 5 |
| Model 8 | Normoxia and anoxia grand maternal effect | mode 1 | mode 2 | mode 3 | mode 4 | mode 5 | mode 6 | mode 7 | mode 8 |

